# 17q21.31 locus regulates Parkinson’s disease relevant pathways through *KANSL1* activity

**DOI:** 10.1101/2025.03.17.641409

**Authors:** Amy R. Hicks, Benjamin O’Callaghan, Jonathan W. Brenton, Melissa Grant-Peters, Aine Fairbrother-Browne, Guillermo Rocamora Perez, Caitlin A. Loh, Regina H. Reynolds, Emil K. Gustavsson, Kylie Montgomery, Maria Tsalenchuk, Raquel Real, Huw Morris, Katie Lunnon, Sarah J. Marzi, Zane Jaunmuktane, John Hardy, Helene Plun-Favreau, Mina Ryten

**Affiliations:** Aligning Science Across Parkinson’s (ASAP) Collaborative Research Network, Chevy Chase, MD, 20815 USA; Department of Genetics and Genomic Medicine, Great Ormond Street Institute of Child Health, University College London, London, UK; NIHR Great Ormond Street Hospital Biomedical Research Centre, University College London, London, UK; UK Dementia Research Institute at The University of Cambridge, Cambridge, UK; Department of Neurodegenerative Disease, UCL Queen Square Institute of Neurology, London, UK; Department of Clinical and Movement Neurosciences, UCL Queen Square Institute of Neurology, London, UK; Queen Square Brain Bank for Neurological Disorders, UCL Queen Square Institute of Neurology, London, UK; UK Dementia Research Institute at University College London, London, UK; Department of Clinical Neuroscience, School of Clinical Medicine, The University of Cambridge, Cambridge, UK; UK Dementia Research Institute, King’s College London, London, UK; Department of Basic and Clinical Neuroscience, Institute of Psychiatry, Psychology and Neuroscience, King’s College London, London, UK; Department of Brain Sciences, Imperial College London, London, UK; Department of Clinical and Biomedical Sciences, Faculty of Health and Life Sciences, University of Exeter, Exeter, UK

## Abstract

An inversion polymorphism at the 17q21.31 locus defines the H1 and H2 haplotypes, with the former linked to multiple neurodegenerative disorders, including an increased risk of Parkinson’s disease (PD). Although the high linkage disequilibrium at this locus has made it difficult to decipher which gene(s) drive the PD association, there is increasing evidence to support the role of *KANSL1* as a risk gene. KANSL1 has been shown to regulate the expression of some PD-associated genes and pathways, likely as part of the histone acetylating non-specific lethal (NSL) complex. Here for the first time, we studied the global effects of 17q21.31 haplotype variation using bulk and single-nuclear RNA-sequencing data from control and PD patient brain. We first analysed differential gene expression across haplotype groups, and then assessed the contribution of *KANSL1* by comparing with the results of an siRNA knockdown in neuronal and glial human cell lines. We demonstrated that the PD risk-associated H1 haplotype downregulates autophagy, lysosomal and mitochondrial processes, all of which have already been implicated in PD aetiology. Furthermore, these effects were apparent in both neuronal and glial cell types, and in the case of the latter, appear to be associated with the modulation of innate and adaptive immune responses. Thus, we identify important links between NSL complex activity and PD pathophysiology that can be leveraged for novel therapeutic interventions.

## Introduction

The 17q21.31 locus is associated with increased risk of multiple neurodegenerative diseases, of which Parkinson’s disease (PD) is the most common[1–4]. First identified as a risk locus in 2001, it has thenceforth been amongst the most strongly-associated PD risk loci in successive studies[5]. However, the molecular mechanisms underlying the link to disease have been challenging to identify and are widely disputed. In part, this is because the locus lies within a linkage disequilibrium (LD) block of approximately 1.5 Mb arising due to a 970 kb inversion polymorphism, and which gives rise to two major haplotypes (termed H1 and H2). The H2 haplotype is most common in Norther European individuals, with much lower frequency in East Asian and African individuals[6]. The H1 haplotype is associated not only with an increased risk of PD, but also the primary tauopathies, Progressive Supranuclear Palsy and Corticobasal Degeneration[7,8]. As such, these risk associations are frequently attributed to *MAPT* due to its role in encoding Microtubule Associated Protein Tau (MAPT). However, PD is not widely considered to be a tauopathy, and although tau and beta-amyloid co-pathology can occur in PD – most commonly in the context of dementia – this form of pathology is not a defining feature of the disease[9].

Inevitably, this has led to increasing interest in the potential contribution of other genes at the 17q21.31 locus to PD risk. In fact, it is known that haplotype variation at the 17q21.31 locus alters the expression of multiple genes therein with a degree of tissue and cellular specificity. Focusing on human brain, there is already evidence to support the divergent expression of *PLEKHM1*, *CRHR1, LRRC37A/2* and *KANSL1* between the two major haplotypes at the tissue-level, with studies of *LRRC37A/2* highlighting a role in astrocytes in particular [10,11]. Nonetheless, we still do not know whether variation at this locus is capable of mediating broader, transcriptome-wide effects on gene expression in human brain and whether such effects could contribute to the regulation of disease-relevant processes in PD. Addressing this key knowledge gap using expression quantitative trait locus (eQTL) analysis, a statistical method that aims to associate genetic variation at a locus with a change in gene expression, is highly challenging, particularly in human brain, due to the very large numbers of samples required to achieve robust results[12]. Indeed, cis-eQTL analysis identifying single nucleotide polymorphisms (SNPs) regulating the expression of 17q21.31 genes (including *KANSL1, PLEKHM1* and *ARL17A)* with varying frequency across eight major cell types required 391 human brains [13]. However, by leveraging existing knowledge of gene function and additional orthogonal forms of analysis, the impact of disease risk variants on global gene expression can be identified, with significant implications for the understanding of the molecular mechanisms driving disease risk.

With these factors in mind, there are compelling reasons to further study the role of *KANSL1* in mediating PD risk at the 17q21.31 locus. Together with the PD GWAS-linked gene *KAT8, KANSL1* encodes components of the Non-Specific Lethal (NSL) complex[14]. This complex consists of nine proteins in total and has a major role in chromatin remodelling, so providing an obvious means for global regulation of gene expression. The effect of perturbing *KANSL1* has been partially characterised, albeit outside the field of neurodegeneration. Haploinsufficiency of *KANSL1* causes Koolen-de Vries syndrome, a rare disorder characterised by symptoms including intellectual disability and epilepsy[15]. Investigations into *KANSL1* loss of function aiming to identify therapeutic strategies for this syndrome have highlighted effects on pathways such as oxidative stress, autophagosome-lysosome fusion and neuronal network activity[16,17]. Furthermore, *KANSL1* has already been implicated in the regulation of important PD pathways, namely mitophagy[18].

Here, we aimed to investigate the regulatory activity of the 17q21.31 locus on global gene expression in human brain, and more specifically the contribution of *KANSL1* (Figure 1). To achieve this, we combined genetically stratified analyses of bulk and single nuclear (sn) RNA-sequencing data derived from post-mortem human brain originating from individuals with PD and neurotypical controls, together with analyses of genetic knockdowns in neuroblastoma and astroglioma cell models. Thus, we tested the hypothesis that the 17q21.31 locus drives differential gene expression that is globally important in the development of PD, and that is mediated at least in part through the gene regulatory activity of the NSL complex.

**Fig. 1.**
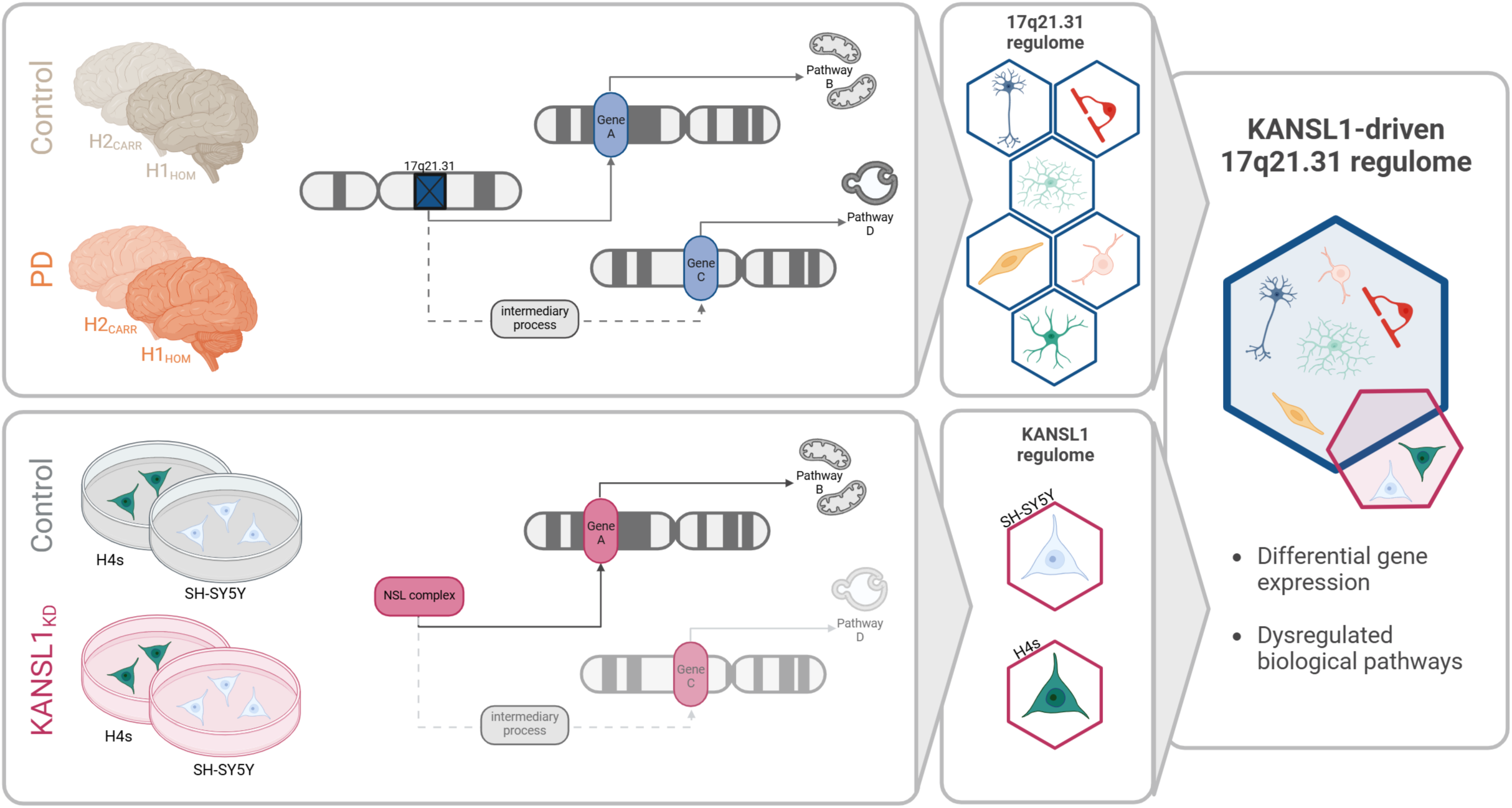
Investigating the regulatory activity of the 17q21.31 locus on global gene expression in human brain, specifically focusing on the contribution of *KANSL1*. We performed the analysis of bulk and snRNA-sequencing data derived from post-mortem human brain tissue originating from individuals with PD and neurotypical control. Samples were stratified according to their 17q21.31 haplotype status, with the goal of characterizing the downstream effects of the regulatory activity of this locus on a cell-type specific level. In parallel, we performed analyses of neuroblastoma and astroglioma cell models with *KANSL1* knockdown to characterise the effects of NSL complex perturbation. Subsequently, in examining common genes and pathways dysregulated in both of these analyses, we assessed the contribution of the gene regulatory activity of KANSL1 and the NSL complex to differential gene expression driven by the 17q21.31 locus.

## Methods

### Brain sample selection

Samples were provided by the Queen Square Brain Bank. We examined samples from 60 donors, each of whom provided samples from four brain regions, namely the anterior cingulate cortex (ACG), inferior parietal lobe (IPL), medial frontal gyrus (MFG) and medial temporal gyrus (MTG). These regions were chosen to analyse the effects of PD pathology at different stages of disease progression, as previously described [19,20]. Donors consisted of 26 females and 34 males and were categorised into 20 controls and 40 PD cases (all at Braak stage 5 or 6). Samples were further categorised according to their 17q21.31 haplotype status, as is summarised in Table 1.

**Table 1).** 17q21.31 haplotype status of all individual donors.

|  | H1/H1 | H1/H2 | H2/H2 |
| --- | --- | --- | --- |
| Control | 13 | 7 | 0 |
| PD | 26 | 13 | 1 |

### Ethics approval

Each tissue donation has informed consent to use the postmortem brain material for research. Tissues were stored and analysed in accordance with UK legislation and the Human Tissue Authority license held by UCL Queen Square Institute of Neurology. The research project was approved by the NHS Health Research Authority, Ethics Committee London-Central.

### Brain sample preparation

As previously described, brain samples were flash-frozen at the time of post-mortem collection and stored at-80°C in the Queen Square Brain Bank, before being dissected on dry ice[20]. All dissections were performed by a single neuropathologist (ZJ) to ensure consistency and pan-cortical representation across samples. Each sample contained approximately 100 mg full-thickness cortex along with overlying leptomeninges.

### RNA isolation for bulk RNA-sequencing from brain

RNA was extracted from brain tissue samples by BioXpedia A/S (Denmark). Samples were first lysed with QIAzol and RNA was extracted using the RNeasy 96 Kit (Qiagen) with an on-membrane DNase treatment, as per the manufacturer’s instructions. Samples were then quantified by absorption on the QIAxpert (Qiagen) and RNA integrity number (RIN) was measured using the Agilent 4200 Tapestation (Agilent).

### Generation of bulk RNA-sequencing from brain

Bulk RNA-sequencing libraries were constructed and sequenced by the UCL Genomics Facility. Total RNA (500 ng) was used as input for the construction of complementary DNA (cDNA) libraries with the KAPA messenger RNA (mRNA) HyperPrep kit (Roche), following manufacturer’s instructions. xGen Dual Index UMI adapters (Integrated DNA Technologies, Inc.) were added to each sample to minimise the mis-assignment of reads when later performing multiplexing. Libraries were multiplexed on S2 and S4 NovaSeq flow cells for paired-end 150 bp sequencing on the NovaSeq 6000 Sequencing System (Illumina), and had a target read depth of 110 million paired-end reads per sample. Finally, sequenced reads were de-multiplexed and BCL Convert software (Illumina) was used to generate FASTQ files.

### Processing of bulk RNA-sequencing data from brain

FASTQ files were processed using a publicly available nextflow pipeline (https://github.com/Jbrenton191/RNAseq_splicing_pipeline.git). Briefly, fastp (v0.23.2) was used for adapter trimming, read filtering and base correction with default settings[21]. Reads with ≥ 40% Phred quality scores below 15, with ≥ 5 bases that could not be called, or shorter than 36 bases were filtered out and adapter sequences were trimmed. Quality control (QC) metrics were visualised prior to alignment using FastQC[22]. Salmon (v 1.9.0) was used to align filtered reads to the transcriptome (Gencode v.41), correcting for sequence-specific, fragment GC content and positional biases and using the entire genome (GRCh38) as a decoy sequence [23,24]. RSeQC (v4.0.0), Qualimap (v2.2.2a) and Picard (v2.27.5) were used for alignment QC. MultiQC (v1.13) was used to visualise QC metrics across all pipeline modules.

Transcriptome-aligned reads generated from Salmon were transformed into gene-level count matrices using the tximport R package (v1.30.0)[25]. Genes overlapping with ENCODE problematic regions, termed the ENCODE blacklist, were removed from downstream analysis[26]. Gene-level expression was filtered to include only genes with count > 0 across each sample group, with 26,207 genes analysed in control samples, 23,403 genes in PD samples and 22,766 genes in all samples grouped together.

### Analysis of bulk RNA-sequencing data from brain

Before analysing differential gene expression, covariates driving variation in the expression data were elucidated using an unbiased, data-driven approach, as previously described[20]. First, all technical, clinical, and histopathological metadata were collated.

Uninformative or unusable variables were removed, including those with zero variance or high missingness. We selected one variable from each pair with high collinearity. Of the variables remaining, those with missing values underwent mean imputation (for numericals), or common value imputation (for categoricals). For assessing pairs of continuous variables, collinearity was assessed using the caret R package (v6.0.94), with a Spearman’s rank correlation coefficient cut-off set at 0.7. For every correlation above this threshold, the variable with the lowest average correlation was kept and the other removed. For assessing pairs of categorical variables, a χ-squared test was used; for pairs of numeric and categorical variables, we used the Kruskal-Wallace test; and for pairs of numeric variables we used a Spearman’s rank correlation test. The resulting *p*-values were assessed using a significance threshold adjusted for multiple comparisons (p < 0.05/(number of tests)). Variables were removed from pairs that demonstrated significant relationships. The contribution to variance in gene expression of the remaining variables was assessed by two methods: 1) variancePartition (v1.35.5), which assesses the contribution of variance of putative covariates at the gene level; and 2) Principal Component Analysis (PCA) dimensionality reduction, which generates per-sample eigenvectors which can then be correlated with putative covariates to assess their relative importance[27]. We employed these two methods to determine our final covariates, recognising the need to control not only for covariates with global impacts on expression, but also those that may confound the expression of small subsets of genes. Covariates were then assessed against three metrics: 1) variancePartition: the maximum variance explained percentage, providing an assessment of whether a covariate comprises a large proportion of the variation of any gene(s); 2) variancePartition: the 3rd quartile of the variance explained distribution, providing an assessment of whether a covariate comprises a sizeable proportion of the variation of a larger proportion of genes; and 3) the Spearman’s rank correlation coefficient between PC-X and covariate. For each metric, data-driven cut-offs were set using visualisation. This resulted in the selection of the following covariates for the differential gene expression model (in addition to a variable representing 17q21.31 haplotype group and disease status): *GC_NC_40:59 + MEDIAN_3PRIME_BIAS + uniquely_mapped_percent + reads_aligned_genes + tss_up_1kb_tag_pct + sex + percent_GC*.

We then performed outlier detection using a method adapted from Gandal et al., 2022 and normalised covariate-corrected expression values[28]. In brief, a sample was considered an outlier if it 1) had an absolute Z-score > 3 for any of the top 10 principal components, 2) had a sample connectivity score <-2, calculated using the WGCNA R package[29].

Differential gene expression analysis was performed using limma (3.46.0) [30,31]. Briefly, the voom function was used to shrink the dispersion and the duplicateCorrelation function was used to remove the effect of each individual, allowing multiple brain regions to be modelled together. These functions were applied twice, as recommended by the developers. To assess the effects of 17q21.31 haplotype status, H2 allele carriers were compared to H1 homozygotes. Haplotype group and brain region were merged into a single term to allow the model to be run just once and allow all comparisons to be extracted. The contrasts were run as a merged analysis incorporating all four brain regions and within three clinical groups: 1) all samples (additionally corrected for disease status); 2) control only; and 3) PD only. Differentially expressed genes (DEGs) were extracted for each contrast using the eBayes and topTable limma functions. False discovery rate (FDR) multiple test correction was applied, with a significance cut-off of < 5 x 10^-2^.

### Generation of single-nuclear RNA sequencing data

Nuclei were isolated from frozen brain tissue using a protocol described here: https://dx.doi.org/10.17504/protocols.io.yxmvm25xng3p/v1. As previously described, a dounce tissue grinder with a custom homogenization buffer was used to homogenize ∼100 mg frozen tissue per sample[20]. Samples were layered onto a density gradient medium and centrifuged using an Optima XPN100 ultracentrifuge at 7700 rotations per minute for 30 minutes at 4◦C in an SW 41 Ti swinging-bucket rotor (Beckman Coulter). The gradient was removed and nuclei were then resuspended and filtered twice to remove further debris. Samples were then counted on a LUNA-FL dual fluorescence cell counter (Logos Biosystems) and diluted to a concentration required to capture 8000 nuclei per sample. Samples were then processed using 10X Genomics GEM isolation technology with the Chromium accessory, and cDNA amplification and library construction was performed using the Single Cell 3’ Reagent Kits, following manufacturer’s instructions. Amplified cDNA and cDNA libraries were then quantified on a QUBit 4 fluorometer (ThermoFisher) and the distribution of molecule sizes was determined using the Agilent 4200 Tapestation (Agilent). Pooled libraries were then loaded onto S4 flow cells and sequenced using the Novaseq 6000 Sequencing System (Illumina).

### Processing of single-nuclear RNA-sequencing data

As previously described, a modified version of the dev branch of the nf-core scrnaseq nextflow pipeline (PMID: 32055031, https://zenodo.org/records/6656322) was used to process snRNA-sequencing data. The v2.0.1 dev (commit: #0104519) was modified to add an option to make matrix conversion optional, which is detailed here https://github.com/RHReynolds/nf-core-scrnaseq/. Read mapping to GRCh38 human reference genome was performed using STARsolo (STAR v2.7.8a) along with gene annotations from Ensembl v107[32–34]. ENCODE standard options for long RNA-sequencing were used, with the exception of (i) –outFilterMultimapNmax (which was set to 1, thus retaining only uniquely mapped reads) and (ii) –alignSJDBoverhangMin (which was set to the STAR default of a minimum 3 bp overhang required for annotated spliced alignment). To generate a filtered gene/cell count matrix almost identical to that of CellRanger, parameters were set as described in STARsolo documentation.

The panpipes package was used to ingest aligned data for QC, preprocessing and clustering, as previously described[20,35]. Briefly, nuclei were filtered according to several QC metrics, namely doublet detection score (score<0.15), percentage of mitochondrial transcripts (<5%), and percentage of ribosomal transcripts (<5%) [36]. Batch correction was performed using Harmony and community detection was performed with the leiden algorithm[37]. Major cell types were detectable with cell type markers (Astrocytes: *AQP4, GFAP*; Endomural cells: *CLDN5*; Neurons: *GABRB2*; Inhibitory neurons: *GAD2*; Immune cells: *CD74, PTPRC, TREM2, APOE*; Oligodendrocytes: *PLP1, MBP*; OPCs: *PDGFRA*, *BCAS1*). Cluster annotations were performed using a combination of cell markers and the highly variable genes calculated for each cluster, implemented by scanpy[38]. Finally, the aggregateToPseudoBulk function from the Dreamlet R package (v0.99.16) was then used to sum gene expression counts of nuclei by donor ID, brain region and cell type annotation[39].

### Analysis of single-nuclear RNA-sequencing data

Again, covariates that were driving variation in gene expression were elucidated using the same approach as described for the bulk RNA-sequencing data[20]. Covariates passing the cut-offs in > 50% cell types were selected as the final covariates, resulting in the selection of the following covariates for the differential gene expression model (in addition to a variable representing 17q21.31 haplotype group and disease status): *sex + RIN* + *deletion_length + insertion_length* + *uniquely_mapped_percent*.

Following pseudobulking, we performed outlier detection on pseudobulked, normalised (log_2_ Fragments Per Million), covariate-corrected expression values using a similar method to that described previously[28]. Again, a sample was considered an outlier if it: 1) had an absolute Z-score greater than three for any of the top 10 principal components, 2) had a sample connectivity score of less than-2, and (3) fulfilled the first two criteria in > 50% cell types. Sample connectivity was calculated using the WGCNA R package[29].

Cell type proportions were compared between H2 allele carriers to H1 homozygotes using the Crumblr R package (https://github.com/GabrielHoffman/crumblr/). Briefly, the frequency of each cell type within each sample was summed, data was covariate corrected using the model described above and the fit was smoothed using eBayes. Analysis was completed within two clinical groups: 1) control only; and 2) PD only. FDR multiple test correction with a significance cut-off of < 5 x 10^-2^ was applied.

Finally, differential gene expression was analysed using the Dreamlet R package[39]. Pseudobulked gene expression counts were first normalised, setting the minimum number of reads per cell type for a gene to be considered expressed to 10. Differential gene expression analysis was then performed using the limma-style set up of Dreamlet’s predecessor, Dream, and comparing H2 allele carriers to H1 homozygotes[40]. Analysis was again completed within the two clinical groups referenced above. FDR multiple test correction with a significance cut-off of < 5 x 10^-2^ was applied.

### Lentivirus generation

Lenti-X 293T human embryonic kidney (HEK) cells sourced from Takara Bio (632180, RRID: CVCL_4401) were cultured in Dulbecco’s Modified Eagle Medium (DMEM, Thermofisher, 11995-065) supplemented with 10% v/v heat inactivated foetal bovine serum (FBS, Thermofisher, A5256801) and maintained in a humidified 37 °C incubator, 95%/5% air/CO2. To generate single guide RNA (sgRNA)-encoding lentiviral particles, 75-95% confluent 6-well dishes of Lenti-X 293T HEK cells were transfected with pMD2.G and pCMVR8.74, alongside sgRNA encoding pLV[Exp]-U6>sgRNA-hPGK>mApple transfer plasmids (Vectorbuilder, Table 2) at a 1:1:2 molar mass ratio using Lipofectamine 3000 (Invitrogen). A full media change was performed 18 hours later, with 2ml of mTeSR Plus (StemCell Technologies) per well. Cells were cultured for 24 hours, at which point the lentivirus-containing mTeSR Plus supernatant was collected and diluted 1:1 with fresh mTeSR Plus supernatant (0.5x), before filtering through 0.44 µm PES filters. pMD2.G (Addgene plasmid #12259, RRID:Addgene_12259) and pCMVR8.74 (Addgene plasmid #22036, RRID:Addgene_22036) were gifts from Didier Trono.

**Table 2).**
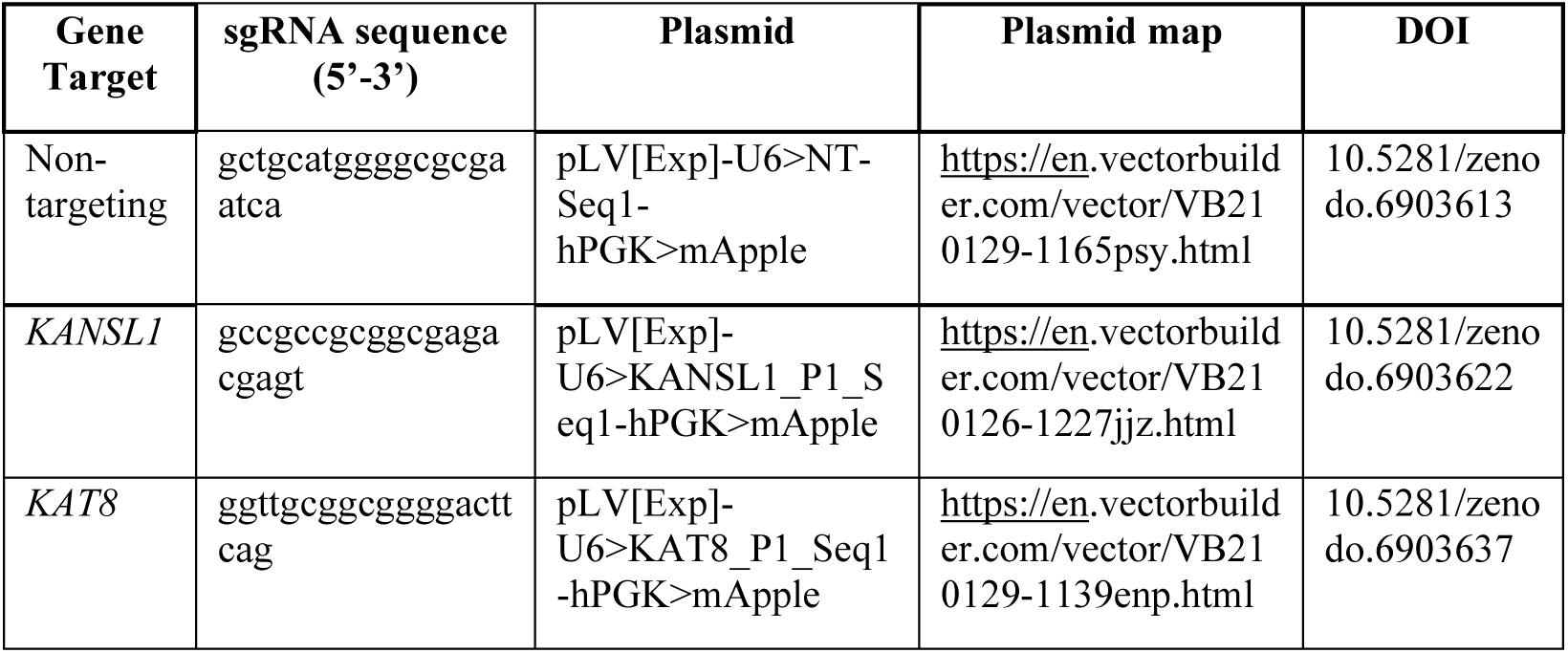
sgRNA sequences and encoding plasmids used in this study.

### Cell culture and treatment

Wildtype SH-SY5Y neuroblastoma cells (RRID:CVCL_0019) and H4 astroglioma cells (RRID:CVCL_1239) were sourced from American Type Culture Collection and cultured in DMEM containing 10% v/v heat inactivated FBS in a humidified incubator at 37°C, as described above. Non-targeting/scrambled (SCR) (D-001206-13), KAT8-targeting (M-014800-00) and KANSL1-targeting (M-031748-00) short interfering RNA (siRNA) were purchased as pre-designed siGENOME SMARTpools and transfected as per the manufacturer’s instructions. Briefly, 4 x 10^5^ SH-SY5Ys or 2.6 x 10^5^ H4s were reverse transfected with 100nM siRNA and 0.5% v/v Dharmafect1 following the manufacturer’s instructions. Following 72 hours incubation, culture media was removed, cells were washed in ice-cold phosphate-buffered saline (PBS) (Gibco, 14190-094) and either centrifuged for 5 minutes at 300 g, 4◦C and stored at-80° as dry pellets, or snap frozen within the plate at-80°.

*KANSL1* and *KAT8* CRISPR interference (CRISPRi) knockdown human iNeurons were generated as previously described[18]. The WTC11 human induced pluripotent stem cell (hiPSC) line which has been incorporated with a doxycycline-inducible Ngn2 system at the AAVS1 locus (iNeuron system) and dead Cas9-KRAB transcriptional repressor fusion protein at the CLYBL locus (CRISPRi system) was used and was a kind gift from the labs of M. E. Ward and M. Kampman[41,42]. hiPSCs were cultured on Geltrex (Thermofisher, A1413202) coated culture dishes in mTeSR Plus medium (StemCell Technologies) and maintained in a humidified 37 °C incubator, 95%/5% air/CO_2_.

*KANSL1* and *KAT8* were knocked down through lentiviral delivery of gene specific sgRNA (see Table 2 for sgRNA sequences) and differentiated into iNeurons through doxycycline induced Ngn2 overexpression. On day-1, hiPSCs were dissociated into a single cell suspension using TrypLE (Thermofisher, 12605010) and reverse transduced with sgRNA lentiviral supernatant in mTeSR Plus supplemented with 5ug/ml polybrene (Sigma, 107689) and 10 µM Y-27632 ROCKi (Apollo Scientific, BISN0135), final 0.25x lentiviral supernatant. In each well of a Geltrex coated 6-well, 4.5 x 10^5^ cells were seeded at a final volume of 1.5ml. iNeuron differentiation was started 24 hours later (day 0) by replacing the media with 2ml of induction media per well, consisting of DMEM/F12 (Thermofisher, 11330032) supplemented with 1x non-essential amino acids (NEAAs, Thermofisher, 11140050), 1x N2-supplement (Thermofisher, 17502048) and 2 µg/ml doxycycline (Sigma, D9891). Full induction media changes were performed on day 1 and day 2. On day 3, the differentiating iNeuron cultures were dissociated using Accutase (Sigma, A6964) and a single cell suspension prepared in N2B27 media consisting of a 1:1 mixture of DMEM/F12:Neurobasal supplemented with 0.5x N2-supplement, 0.5x B27 supplement (Thermofisher, 17504044), 0.5x NEAAs, 0.5x Glutamax, 45 µM 2-Mercaptoethanol (Thermofisher, 21985023) and 2.5 µg/ml insulin (Sigma, I9278). Following this, 6 x 10^5^ cells were seeded per 12-well in 1ml final volume. A half media change was performed with N2B27 24 hours later (day 4) and every 3-4 days thereafter. A half media change with N2B27 was performed on day 16, 24 hours prior to collection for RNA extraction on day 17.

### RNA isolation from cell lines

Total RNA was extracted from dry cell line pellets and iNeurons following the manufacturer’s protocol for the Monarch Total RNA Miniprep Kit (New England Bioscience, T2010) with inclusion of the optional on-column DNAse treatment and quantified using a NanoDrop One Spectrophotometer (Thermofisher Scientific). Cell line RNA extracted for sequencing was diluted to 50 ng/μL in RNase-free water.

### Quantitative real time PCR

iNeuron RNA was reverse transcribed in a 10 µl reaction with 2.5 U/µl SuperScript IV reverse transcriptase (Thermofisher, 18090050) with 2.5 µM random hexamers (Thermofisher, SO142), 0.5 mM dNTPs (Thermofisher, R0191), 5 mM DTT and 2 U/µl RNAseOUT (0777019). Equal amounts of RNA were reverse transcribed for all samples in a single experiment, with 200 ng RNA in a 10 µl reverse transcription reaction being the most common. The cDNA product was diluted such that 200ng reverse transcribed RNA would be in an 800ul final volume (i.e. 0.25 ng/µl). Next, 4 µl diluted cDNA (i.e. 1 ng) was subjected to quantitative real-time PCR (RT-qPCR) using 1x Fast SYBR™ Green Master Mix (Applied Biosystems) and 500 nM gene specific primer pairs (Table 3) on a QuantStudio™ 7 Flex Real-Time PCR System (Applied Biosystems). At least three technical replicates were performed for each sample and gene target combination, and a RT-control for all samples and gene target combinations was performed alongside. Relative mRNA expression levels were calculated using the 2−ΔΔCt method with *UBC* as the stable house-keeping gene. *KANSL1* knockdowns were compared to SCR controls using a paired two-tailed Student’s T-test.

**Table 3).**
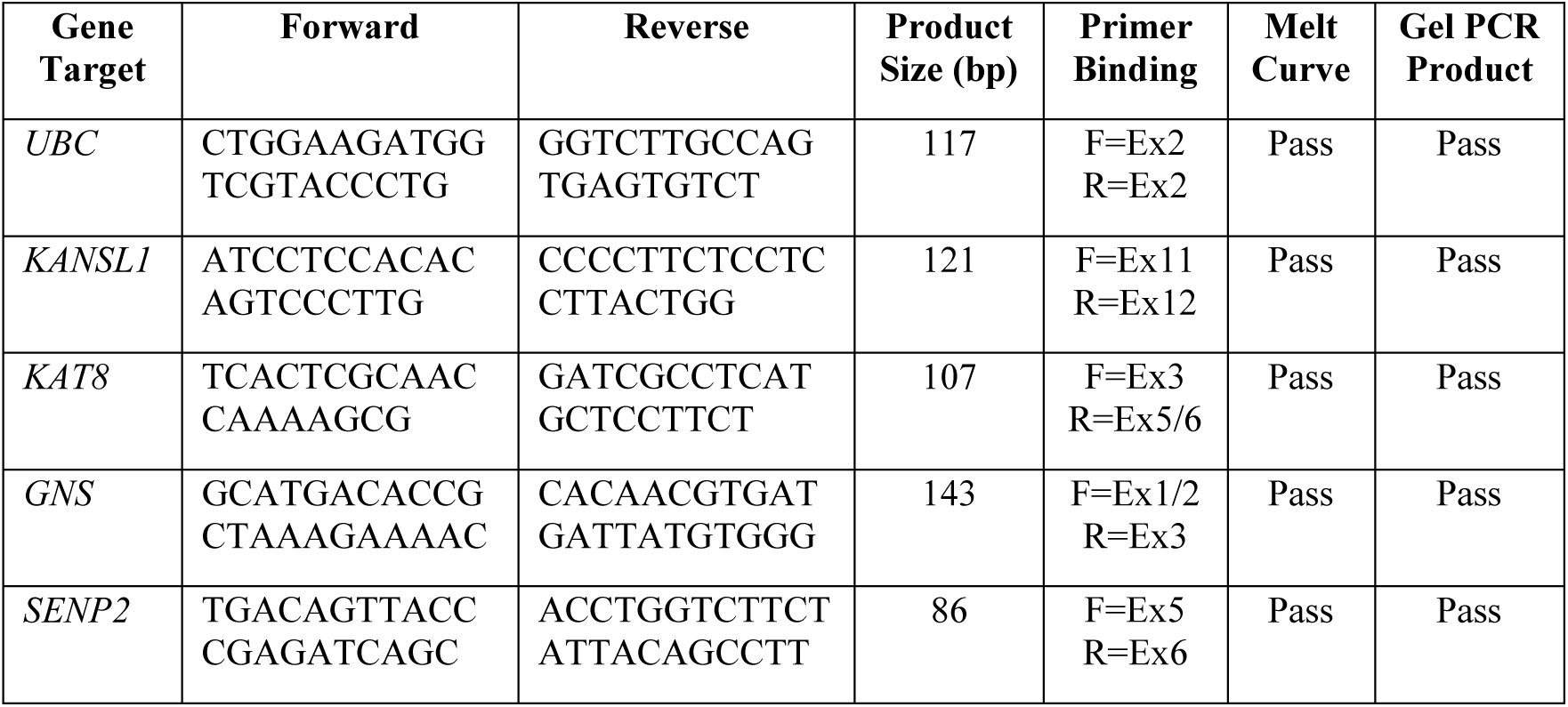
Gene specific primers used for RT-qPCR. Exon (Ex), forward (F), reverse (R).

### Generation of bulk RNA-sequencing data from cell lines

In total, 22 SH-SY5Y samples and 18 H4 samples were sequenced, with the former comprising two separate batches (Table 4). Each batch consisted of multiple biological replicates each of SCR, KAT8-and KANSL1-targetting siRNA-treated cells.

**Table 4).** RNA-sequencing batches of SH-SY5Y and H4 cell lines.

| Batch | Cell type | Number of samples |
| --- | --- | --- |
| Batch 1 | SH-SY5Y | 4 SCR, 2 KANSL1, 4 KAT8 |
| Batch 2 | SH-SY5Y | 4 SCR, 4 KANSL1, 4 KAT8 |
| Batch 3 | H4 | 6 SCR, 6 KANSL1, 6 KAT8 |

Bulk RNA-sequencing libraries were constructed and sequenced by the UCL Genomics Facility using a similar protocol to that which was used for the brain samples, with the following exceptions: libraries were multiplexed on the NovaSeq SP v1.5 for paired-end 100 bp sequencing on the NovaSeq 6000 Sequencing System (Illumina), aiming for a mean read depth of 66 million paired-end reads per sample.

### Processing of bulk RNA-sequencing data from cell lines

Data was processed using the aforementioned nextflow pipeline, accessible here: https://github.com/Jbrenton191/RNAseq_splicing_pipeline.git. Genes overlapping with the ENCODE blacklist were removed from downstream analysis[26]. Gene-level expression was again filtered to include only genes with count > 0 across each siRNA treatment, after which 20,638 genes were detected. The DESeq2 (v1.30.1) R package was utilised to assess differential gene expression with default parameters[43]. Pairwise comparisons were applied across the siRNA treatment groups. A Wald test was used to test for significant differences, wherein the null hypothesis denoted no difference in gene expression. The model consisted of replicate and siRNA treatment. Multiple test correction was performed using FDR correction, with a significance cut-off of < 5 x 10^-2^.

### Generation of gene lists and pathways of interest

Several lists of genes were collated to examine within these RNA-sequencing datasets. The nine genes encoding components of the NSL complex are published[14]. A list of genes associated with Mendelian PD was assembled as previously described[20], combining those in which Blauwendraat et al. (2020) had “High” or “Very High” confidence in them as a Mendelian PD gene, along with searching “Parkinson’s disease” on OMIM[44]. The following genes were manually removed from the latter list due to being disproved or having tenuous evidence supporting PD causation: *GIGYF2*, *UCHL1*, and *EIF4G1*. A list of genes associated with sporadic PD was assembled by combining the results of the Nalls et al., 2019 meta-analysis with that of the Kim et al. 2023 multi-ancestry GWAS[2,45]. In addition, a list of 435 genes associated with the lysosome was obtained from The Human Lysosome Gene Database, which was compiled by collating information from 16 different sources including published proteomic studies and reviews, as well as databases such as gene ontology (GO), Kyoto Encyclopaedia of Genes and Genomes (KEGG), Reactome and UniProt[46]. Finally, we utilised a collection of 46 gene sets genetically implicated in PD through common genetic variation and polygenic risk scoring[47]. This list contained three Pathway Interaction Database (PID) terms, eight KEGG terms and 35 Reactome terms.

### Gene set enrichment analysis

To analyse the enrichment of selected gene sets, *p*-values were obtained using Fisher’s exact tests comparing the overlap in input genes and genes identified as differentially expressed. FDR corrections were used to account for multiple testing where needed. DEGs were functionally annotated with gene ontology (GO) terms, as well as KEGG and Reactome terms, using the gProfiler2 (v0.2.0) R package[48]. GO terms were reduced to parent terms using semantic similarity with varying cut-offs (recorded in figure legends) using the Rutils GitHub package (https://github.com/RHReynolds/rutils) to aid visualisation. In addition, ranked gene set enrichment analysis (GSEA) was performed on results of the differential expression comparing 17q21.31 haplotype effects. Ranked GSEA of GO terms was performed using the gseGO function within the clusterProfiler (v4.8.3) R package, whilst *KANSL1* DEGs, Mendelian and sporadic PD-associated gene lists, and PD-linked KEGG and Reactome pathways were analysed using the fgsea (v1.26.0) R package[49,50]. Analyses with the latter were run with 10,000 permutations. A positive enrichment score (ES) denoted an association with genes with most positive log_2_ fold change (log_2_FC), described as upregulated gene expression, and vice versa.

## Results

### The 17q21.31 locus is associated with increased expression of immune-relevant genes and pathways

To investigate the impact of haplotype variation at the 17q12.13 locus, we used bulk RNA-sequencing data generated from post-mortem human brain tissue originating from 40 PD patients (Braak stage 5/6) and 20 controls unaffected by neurodegenerative disease (Fig. 1). Clinical notes and neuropathology were reviewed to ensure diagnostic accuracy of all donors, as previously described[20]. The mean age at death across clinical groups ranged from 79 years in PD to 87 years old in controls, with slightly higher male:female ratios for PD (25:15) than for controls (9:11), reflecting the gender differences observed in PD. For each individual, four brain regions were sampled, namely the anterior cingulate gyrus (ACG), inferior parietal lobe (IPL), medial frontal gyrus (MFG) and medial temporal gyrus (MTG).

Given the relatively small sample numbers, gene expression values were collapsed across regions to generate a robust measure of expression for each individual. We analysed differential gene expression within three clinical groups, namely controls alone, PD alone, and all samples together, in each case comparing individuals carrying the protective H2 allele (heterozygotes and homozygotes, n = 21) to individuals homozygous for the H1 PD risk allele (n = 39). Interestingly, the greatest number of differentially expressed genes (DEGs) was detected in the control group (12 upregulated, 13 downregulated) (Fig.2a), and across all analyses we found that the majority of DEGs were upregulated in H2 allele carriers compared to H1 homozygotes. As has been demonstrated previously, many genes found to be differentially expressed were located within the 17q21.31 locus itself (Fig. 2b). Elevated expression of six genes therein was observed across every clinical group (FDR range = 2.77 x 10^-39^ – 8.46 x 10^-3^, log_2_ fold change (FC) range = 0.234 – 5.63), whilst elevated *KANSL1* was detected only when control and PD samples were combined (FDR = 8.46 x 10^−3^, log_2_FC = 0.234). Conversely, robust increases in its antisense gene, *KANSL1-AS1*, were detected across all conditions (FDR range = 1.15 x 10^-20^ – 1.20 x 10^-4^, log_2_FC range = 1.17 – 1.33). Notably, *MAPT* was not found to be significantly differentially expressed.

**Fig. 2.**
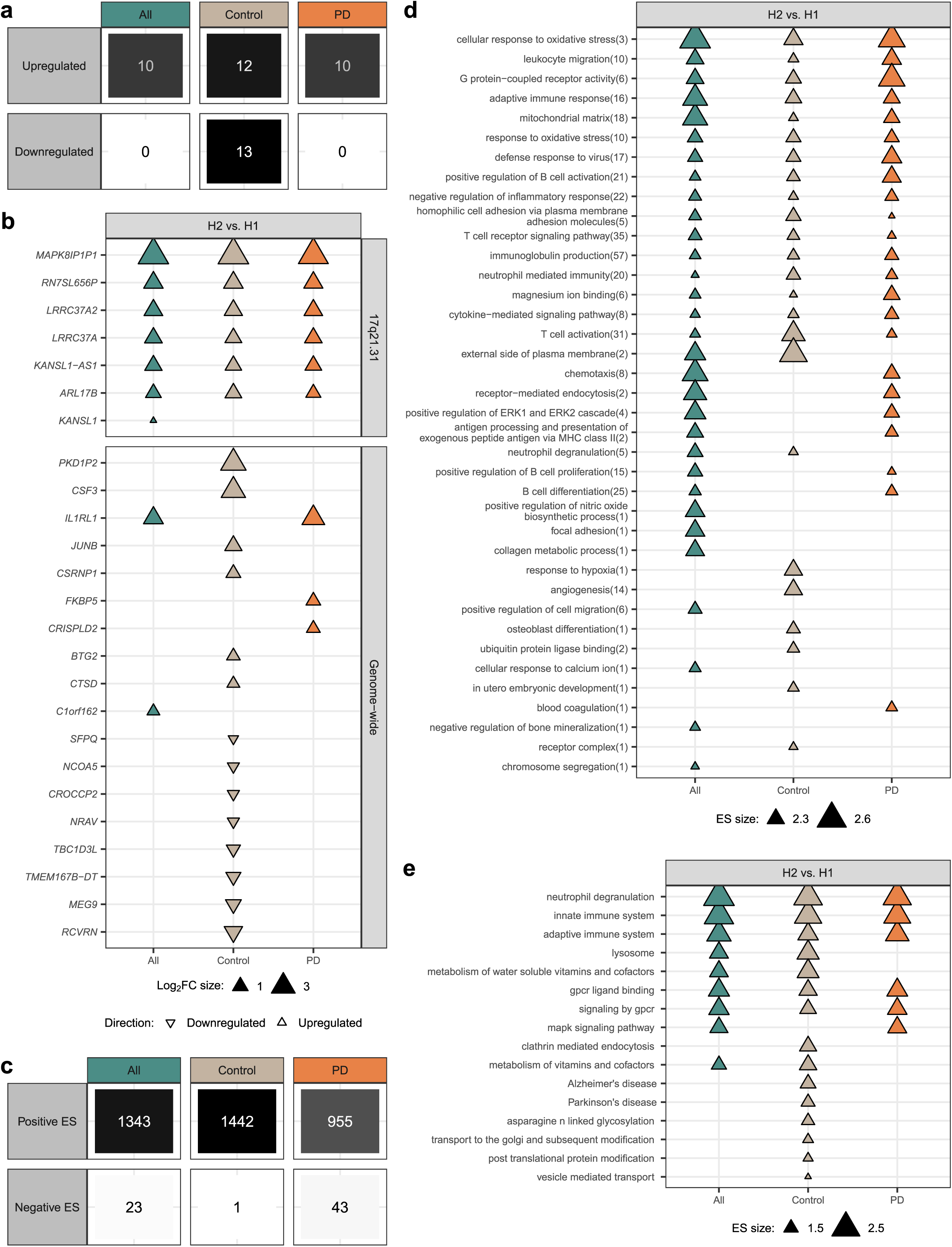
Differential gene expression driven by the 17q21.31 locus is associated with immune system pathways. A) Across three clinical groups, the highest number of DEGs detected when comparing H2 allele carriers to H1 homozygotes was identified in controls. B) Differential expression was detected of genes within the 17q21.31 locus as well as genome-wide, several of which were related to the immune system. C) Using ranked GSEA, we detected the enrichment of GO pathways across differential gene expression in all three clinical groups, particularly among upregulated gene expression. Enrichment score (ES) denotes the strength of the pathway enrichment as well as the direction within the ranked results. D) The top 10% of GO terms enriched across changing gene expression contained many immune system-related pathways. After removing child terms that were enriched in both directions within the same clinical group, terms were ranked according to ES and the top 10% displayed as parent terms, with the number of reduced child terms displayed in brackets (term reduction utilised a hierarchical tree cut-off threshold of 0.9). Triangle size corresponds to the ES of the child term with the least significant FDR. E) Pathways genetically-linked to PD through polygenic risk scoring were also found to be enriched across changing gene expression using ranked GSEA[47].

Importantly, this analysis also identified genes outside the 17q21.31 locus that were differentially expressed in one or more clinical group (Fig. 2b). The highest numbers of such genes were detected in the control group and included *CTSD*, which was upregulated in H2 carriers compared to H1 homozygotes (FDR = 1.80 x 10^−2^, log_2_FC = 0.618). Given that *CTSD* encodes cathepsin D, a protein with a major role in α-synuclein degradation and in the cleavage of prosaposin, an activator of *GBA1*-encoded GCase protein, this finding was of particular interest[51]. We detected additional DEGs through the inclusion of PD samples. More specifically, we noted that *IL1RL1,* which encodes an interleukin 1 receptor protein induced by proinflammatory stimuli, was upregulated in H2 carriers, with the highest effect size seen in PD samples (FDR = 6.35 x 10^-7^, log_2_FC = 2.69)[52]. Similarly, *FKBP5*, which encodes an inflammatory mediator, was upregulated in PD samples alone (FDR = 2.40 x 10^-3^, log_2_FC = 0.863), as was *CRISPLD2* (FDR = 4.78 x 10^−2^, log_2_FC = 0.895), which encodes a protein thought to attenuate excessive immune responses[53,54].

To investigate the possibility that genetic variation at the 17q21.31 locus could impact on pathways including immune responses in human brain, we utilised ranked gene set enrichment analysis (GSEA), ranking by log_2_FC. We identified a total of 1366 GO terms that were significantly associated with changing gene expression in all samples (ES range =-2.09 – 2.82), 1443 in control (ES range =-1.70 – 2.64) and 998 in PD (ES range =-1.99 – 2.58), with the majority of pathways enriched among genes being upregulated in H2 allele carriers (Fig. 2c, Supplementary Fig. 1). Consistent with our findings demonstrating differential expression of genes implicated in immune responses, we noted that focusing on pathways with the highest enrichment scores (top 10% of 3807 total enrichments) highlighted immune activation and regulation (Fig. 2d). These enriched terms included T cell activation (GO: 0042110, FDR range = 4.18 x 10^-11^ – 3.01 x 10^-5^, ES range = 2.23 – 2.28), B cell differentiation (GO: 0030183, FDR range = 5.19 x 10^-7^ – 6.57 x 10^-5^, ES = 2.24) and neutrophil mediated immunity (GO:0002446, FDR range = 1.51 x 10^-27^ – 1.36 x 10^-6^, ES range = 2.28 – 2.51). We also found terms relating to mitochondrial function such as mitochondrial matrix (GO:0005759, FDR range = 9.16 x 10^-14^ – 4.41 x 10^-5^, ES range = 2.22 – 2.47) and response to oxidative stress (GO:0006979, FDR range = 1.67 x 10^-6^ – 2.38 x 10^-5^, ES range = 2.23 – 2.27) were highly enriched. Importantly, these pathway enrichments were highly similar across the control as well as the PD clinical group, suggesting that such global effects on gene expression precede disease development, rather than simply being a consequence of established PD.

### Parkinson’s disease-associated genes and pathways are dysregulated by 17q21.31 haplotype variation across disease states

To assess the relevance of the genes differentially regulated by haplotype status to PD, we collated lists of genes causally linked to Mendelian and sporadic forms of PD[3,44,45]. No gene within either list was differentially expressed between H2 allele carriers and H1 homozygotes in either sample group. We subsequently examined the association of both gene lists across our using ranked GSEA, and found the sporadic PD gene list was nominally enriched amongst genes upregulated in H2 carriers compared to H1 homozygotes in PD samples (FDR = 5.21 x 10^-2^). This enrichment was driven by genes including *NOD2*, *MSR1* and *HLA-DRB5* (log_2_FC range = 0.832 – 0.910) and suggested that downregulation of these genes in H1 homozygotes may be associated with the increased risk of PD previously found at the 17q21.31 locus.

To further explore this association, we utilised a list of 46 pathways genetically implicated in PD based on polygenic risk scoring[47]. We found that 16 terms (34.8%) were associated with upregulated gene expression in H2 carriers compared to H1 homozygotes (Fig. 2e). Interestingly, 15 terms (FDR range = 1.53 x 10^-9^ – 4.22 x 10^-2^) were detected through the analysis of the control group, whilst the inclusion of PD samples, either alone or together with the control group, resulted in the detection of just six terms (FDR range = 1.53 x 10^-9^ – 2.15 x 10^-3^) and nine terms (FDR range = 1.53 x 10^-9^ – 1.52 x 10^-2^) respectively. Enriched terms included the lysosome (KEGG:04142, FDR range = 2.98 x 10^-5^ – 1.52 x 10^-2^) as well as pathways relating to the immune system, namely neutrophil degranulation (R-HSA-6798695, FDR = 1.53 x 10^-9^), innate immune system (R-HSA-168249, FDR = 1.53 x 10^-9^) and adaptive immune system (R-HSA-1280218, FDR = 1.53 x 10^-9^). These results indicate that pathologically relevant pathways, in particular those relating to the lysosome and the immune system, are altered by haplotype variation at the 17q21.31 locus.

### 17q21.31 haplotype variation causes variable differential gene expression across cell types

Next, we examined the effect of 17q21.31 haplotype variation on cell type-specific differential gene expression using snRNA-sequencing data generated from post-mortem human brain tissue from the same 40 PD patients and 20 neurotypical controls (Fig. 1). Two brain regions, namely ACG and IPL, were sampled. All samples underwent nuclear isolation and snRNA-sequencing, and following removal of doublets and low quality cells as previously described, a total of 621,428 nuclei remained in the cohort[20]. Following batch correction, dimensionality reduction and Leiden community detection, nuclei were divided into seven major cell classes: astrocytes, endomural cells, excitatory neurons, immune cells, inhibitory neurons, oligodendrocytes and oligodendrocyte precursor cells (OPCs). Importantly, we detected no significant effects of the haplotype on the proportion of any cell type among either control or PD samples (FDR range = 2.98 x 10^-1^ – 9.75 x 10^-1^).

Using the statistically robust analytical framework provided by the Dreamlet R package, we investigated gene expression by cell type across two clinical groups, namely controls and PD patients[39]. We compared individuals carrying the protective H2 allele to individuals homozygous for the H1 PD risk allele to identify DEGs within each cell type. Overall, we found that control samples harboured greater numbers of DEGs across most cell types (Fig. 3a, Supplementary Fig. 2a). Across all cell types, we detected 710 unique DEGs in total in controls (77.1% in IPL, 30.9% in ACG) and 157 in PD samples (42.7% in IPL, 65.6% in ACG). We noted that most differential gene expression was specific to individual cell types and included both glial and neuronal cell types: 91.8% and 91.0% of DEGs were cell type-specific in IPL control and PD samples respectively; 92.7% and 92.2% of DEGs were cell type-specific in ACG control and PD samples respectively (Fig. 3b,c, Supplementary Fig. 2b,c). The overall lack of overlap between DEGs across cell types points towards the possibility that the gene regulatory activity on the 17q21.31 locus may have significant cell type biases.

**Fig. 3.**
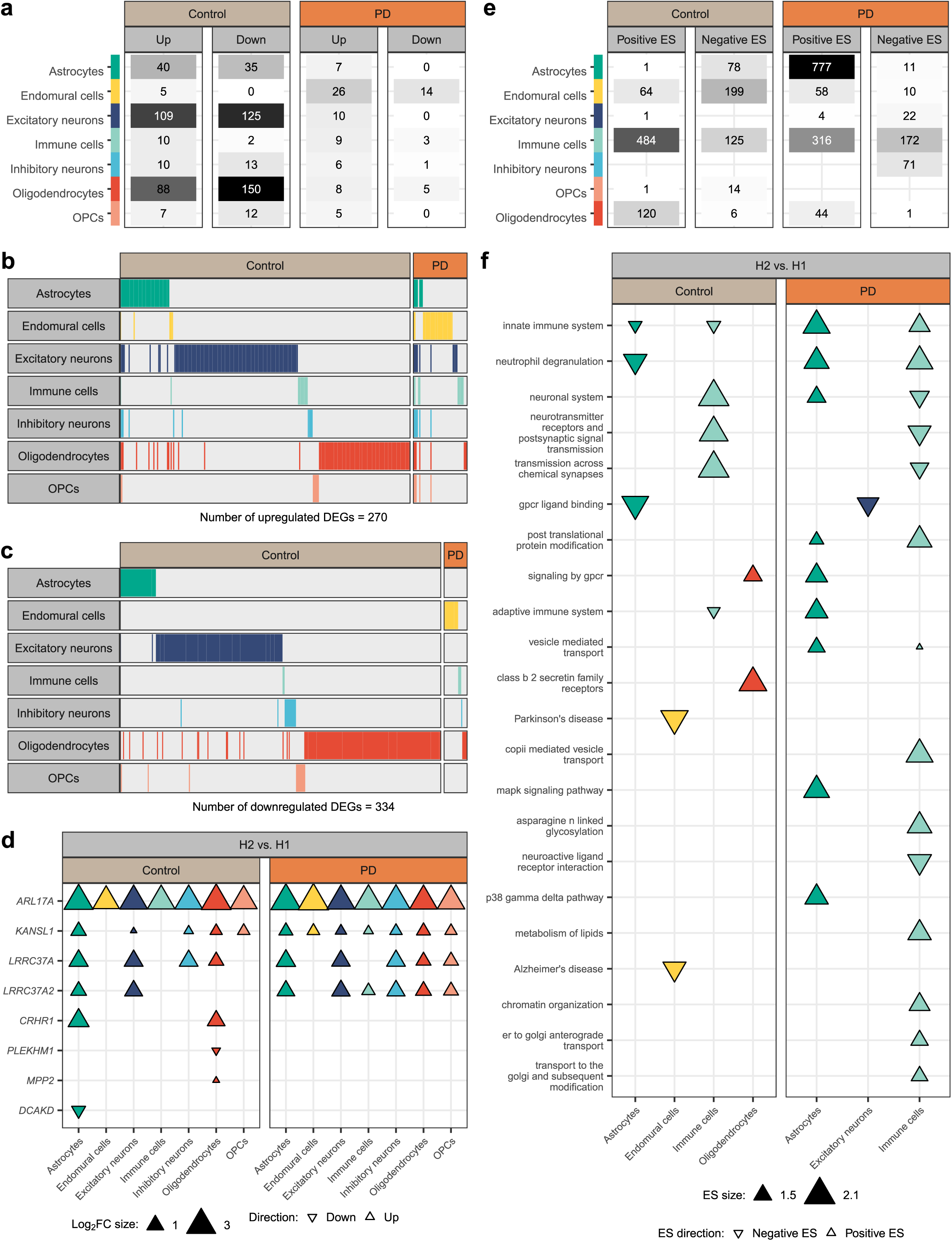
The 17q21.31 locus drives cell type-specific patterns of differential gene expression in IPL. A) Across most major cell types, more DEGs were detected in control than PD samples when comparing H2 allele carriers to H1 homozygotes. B) The majority of differential gene expression detected in this analysis were cell type-specific, in both upregulated and C) downregulated directions. D) We detected differential expression of eight genes within the 17q21.31 locus, three of which were specific to one cell type in one clinical group, with the remaining five dysregulated across multiple cell types. E) Using ranked GSEA, we detected the enrichment of GO pathways across differential gene expression in all major cell types, particularly in astrocytes and immune cells. Enrichment score (ES) denotes the strength of the pathway enrichment as well as the direction within the ranked results. F) Pathways genetically-linked to PD through polygenic risk scoring were also found to be enriched across changing gene expression in four major cell types using ranked GSEA[47].

As expected, many DEGs identified were located within the 17q21.31 locus and were amongst those detected across multiple cell types (Fig. 3d, Supplementary Fig. 2d). We detected the differential expression of a total of 11 genes therein (72.7% differentially expressed in IPL, 90.9% in ACG). *ARL17A* was consistently upregulated in H2 allele carriers compared to H1 homozygotes across all cell types in both control and PD IPL samples (FDR range = 2.47 x 10^-18^ – 1.05 x 10^-2^, log_2_FC range = 2.47 – 3.36) and ACG samples (FDR range = 8.22 x 10^-17^ – 5.51 x 10^-5^, log_2_FC range = 2.39 – 3.17). Similarly, *KANSL1* was found to be upregulated across all cell types in IPL (FDR range = 9.37 x 10^-7^ – 1.36 x 10^-2^, log_2_FC range = 0.527 – 0.755) and ACG PD samples (FDR range = 1.71 x 10^-6^ - 5.65 x 10^-4^, log_2_FC range = 0.555 – 0.850), whilst reaching significance in five cell types in IPL control samples (FDR range = 9.54 x 10^-3^ – 3.18 x 10^-2^, log2FC range = 0.468 – 0.878) and one in ACG control samples (FDR = 4.83 x 10^-2^, log_2_FC = 0.557). This highlights that the differential expression of *KANSL1* is highly consistent across cell types. We compared these results to a published cis-eQTL analysis and found that, of the 11 locus genes that we found to be differentially expressed between haplotype groups, nine had cis-eQTLs in the same cell types[13]. Given the high expression of *MAPT* in neurons, as well as the detection of a cis-regulating SNP in astrocytes by Haglund et al. (2025), we may have expected *MAPT* to be differentially expressed. However, it did not reach significance in any cell type in our analysis of either brain region (FDR range = 2.20 x 10^-1^ – 9.95 x 10^-1^).

We then sought to functionally assess the genes differentially regulated between the haplotype groups across all seven major cell types. Across both clinical groups, ranked GSEA revealed GO terms associated with both upregulated and downregulated gene expression in H2 allele carriers compared to H1 homozygotes (Fig. 3e, Supplementary Fig. 2e). IPL PD astrocytes displayed the largest numbers of terms associated with upregulated gene expression (FDR range = 3.39 x 10^-7^ – 4.94 x 10^-2^) (Fig. 3a). Several of the top terms enriched at the bulk tissue level were also detected at the cellular level in both glial and neuronal cell types (Figure 2d, Supplementary Fig. 3, Supplementary Fig. 4). Common terms such as adaptive immune response (4/7 cell types, GO:0002250, FDR range = 2.52 x 10^-2^ – 4.18 x 10^-2^, ES range =-1.99 – 1.51) and positive regulation of B cell activation (2/7 cell types, GO:0050871, FDR range = 2.69 x 10^-2^ – 4.21 x 10^-2^, ES range = 1.38-1.39) were enriched in IPL, whilst ACG was enriched for terms including cytokine-mediated signalling pathway (5/7 cell types, GO:0019221, FDR range = 2.82 x 10^-2^ – 4.79 x 10^-2^, ES range =-1.66 – 1.56). We also noted terms relating to mitochondrial function, such as mitochondrial inner membrane (6/7 cell types, GO:0005743, FDR range = 4.09 x 10^-4^ – 4.67 x 10^-2^, ES range =-1.99 – 1.76) in IPL and mitochondrial respiratory chain complex I assembly in ACG (4/7 cell types, GO:0032981, FDR range = 5.54 x 10^-4^ – 1.82 x 10^-2^, ES range =-1.73 – 1.45). These results show that important immune-related pathways, as well as mitochondrial mechanisms, were dysregulated by 17q21.31 haplotype variation across a range of major cell types.

### Parkinson’s disease-associated genes and pathways are dysregulated by 17q21.31 haplotype variation in specific cell types

To assess the relevance of DEGs detected between H2 allele carriers and H1 homozygotes across major cell types, we focused on genes associated with Mendelian and sporadic forms of PD[2,44,45]. After excluding genes within the 17q21.31 locus, we found that five genes causally-associated with PD were differentially expressed between H2 carriers and H1 homozygotes in IPL control samples across two cell types: *PRKN, LZTS3, CC2D2A* and *PIK3CA-DT* in oligodendrocytes (FDR range = 6.64 x 10^-3^ – 4.41 x 10^-2^, log_2_FC range =-0.887 – 0.544); and *PMVK* in excitatory neurons (FDR = 2.20 x 10^-2^, log_2_FC =-0.429). An additional PD-associated gene was upregulated in astrocytes and inhibitory neurons of the ACG, namely *DDRGK1* (FDR range = 3.57 x 10^-2^ – 3.62 x 10^-2^, log_2_FC range = 0.412 – 0.931). Furthermore, we used ranked GSEA to examine the enrichment of PD-associated gene lists in each cell type. This revealed that sporadic PD-related genes were upregulated in H2 carriers compared to H1 homozygotes in IPL PD oligodendrocytes (FDR = 7.56 x 10^-3^), potentially indicating that these cells may drive similar pattern detected at the bulk tissue level.

Next, we examined the enrichment of the aforementioned list of 46 gene sets genetically implicated in PD risk across ranked results for each cell type[47]. In total, we found that 22 (47.8%) terms were enriched to varying degrees across changing gene expression in IPL, of which 11 were also enriched in ACG (Fig. 3f, Supplementary Fig. 2f). Overall, more terms were found to be associated with upregulated gene expression in H2 allele carriers compared to H1 homozygotes than downregulated expression. Interestingly, in IPL we found that downregulated gene expression in control glia (astrocytes and immune cells) was associated with terms such as innate immune system (R-HSA-168249, FDR range = 1.56 x 10^-10^ – 6.07 x 10^-5^) and neutrophil degranulation (R-HSA-6798695, FDR = 7.90 x 10^-4^), contrasting to the association with upregulated expression in PD samples (FDR range = 9.19 x 10^-4^ – 1.26 x 10^-2^ and 2.17 x 10^-6^ – 3.13 x 10^-5^ respectively). In summary and combined with the results at the bulk tissue level, these findings suggest that immune related pathways are important in driving the differences in PD risk between 17q21.31 haplotype carriers, and may operate predominantly through glial cell types.

### NSL complex knockdown causes widespread, overlapping differential gene expression

Whilst we expected to detect the differential expression of 17q21.31 locus genes between haplotypes, this raised the question as to which of these genes had the potential to affect the gene regulatory activity of the locus in brain. To further explore this impact, we used *in vitro* systems which enabled us to investigate the role of *KANSL1* in isolation from other genes at the locus and without the issue of secondary effects of disease inherent with studying post mortem tissue. To model the risk-associated H1 haplotype and the accompanying reduction in *KANSL1* expression as compared to H2, we characterised the effects of *KANSL1* knockdown in two immortalised human cell lines of neuronal (SH-SY5Ys) and glial type (H4s).

siRNA knockdown resulted in a 29.2% reduction in *KANSL1* expression on average in SH-SY5Ys (FDR = 3.93 x 10^−4^, log_2_FC = - 0.498) and a 52.0% reduction in H4s (FDR = 1.04 x 10^−43^, log_2_FC =-1.06) (Supplementary Fig. 5a). Furthermore, we detected the dysregulation of 1840 genes in total in SH-SY5Ys (FDR range = 1.90 x 10^−21^ – 4.998 x 10^−2^, log_2_FC range =-4.39 – 3.17), and 7354 genes in H4s (FDR range = 1.07 x 10^−88^ – 4.998 x 10^−2^, log_2_FC range =-5.86 – 4.58) (Fig. 4a).

**Fig. 4.**
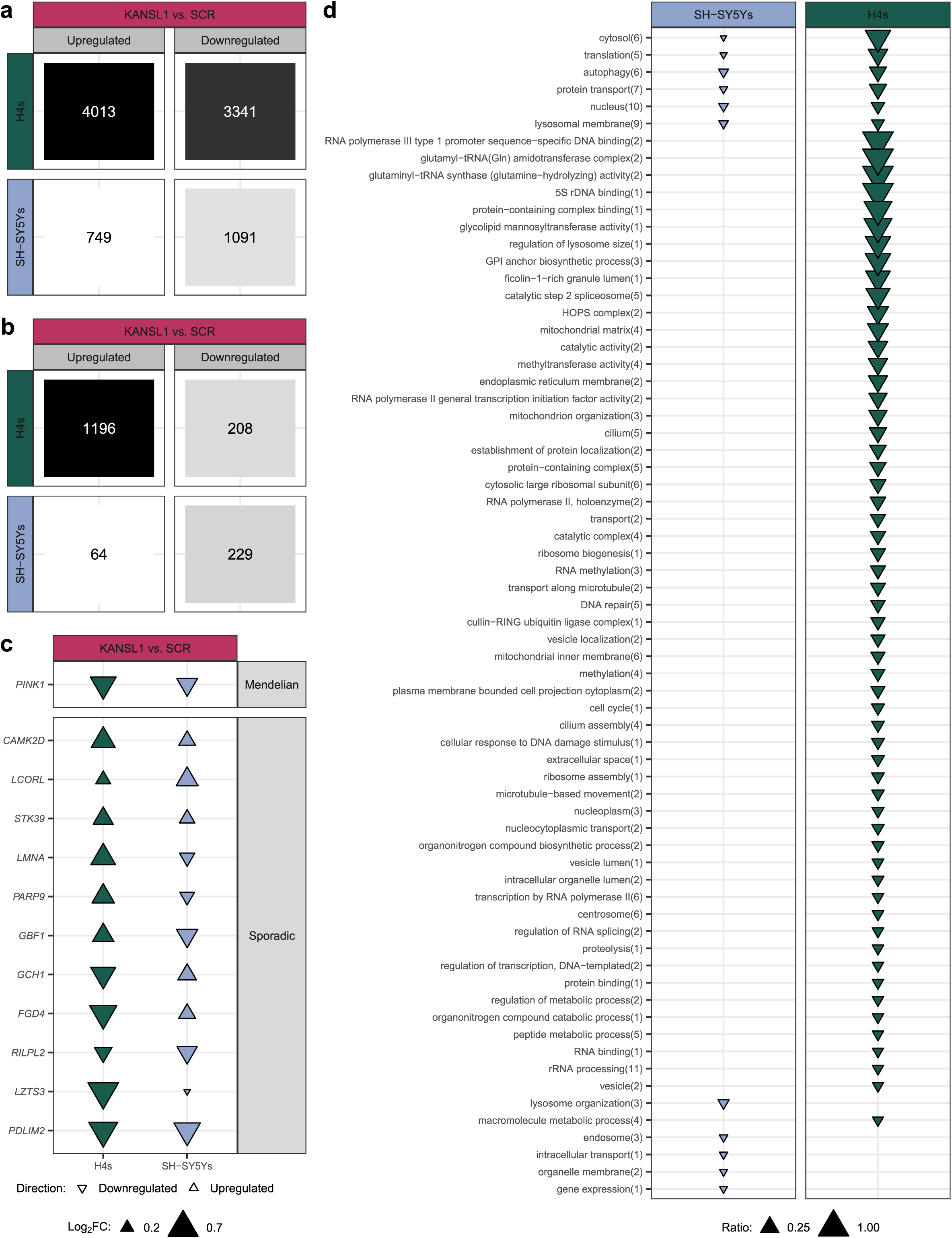
Differential gene expression detected following *KANSL1* knockdown in SH-SY5Y and H4 cells included genes and pathways implicated in PD. A) High numbers of DEGs were detected following *KANSL1* knockdown in both cell lines. B) We detected the enrichment of high numbers of GO pathways among DEGs detected in both cell lines. C) Across both cell lines, we detected differential expression of genes linked to both sporadic and Mendelian forms of PD [3,44,45]. D) Amongst genes that were downregulated following *KANSL1* knockdown and had additional evidence of KANSL1 binding through ChIP-sequencing [16], we detected the enrichment of GO pathways relevant to PD. Parent terms are displayed, with the number of reduced child terms displayed in brackets (term reduction utilised a hierarchical tree cut-off threshold of 0.6). Triangle size corresponds to the ratio (intersection size/term size) of the child term with the least significant FDR.

Since KANSL1 protein acts primarily within the NSL complex, we also investigated the impact of knocking down another component of the complex, namely the established PD risk gene *KAT8*, so enabling us to ensure that our findings could be reliably attributed to NSL functional defects[14]. Although *KAT8* knockdown resulted in the detection of fewer DEGs, with 603 genes dysregulated in SH-SY5Ys (FDR range = 5.54 x 10^−15^ – 4.997 x 10^−2^, log_2_FC range =-2.45 – 3.47) and 5714 genes in H4s (FDR range = 7.96 x 10^−87^ – 4.999 x 10^−2^, log_2_FC range =-3.86 – 3.99), there was high commonality across the gene sets (Supplementary Fig. 5b). We found that *KANSL1* and *KAT8* knockdown DEGs significantly overlapped across both cell types (*p*-value range = 2 x 10^−16^ – 1.00 x 10^-13^, accounting for direction of effect). Furthermore, when studying the DEGs identified through either knockdown, we found highly significant correlations in both cell lines (*p-*value < 2.2 x 10^-16^) (Supplementary Fig. 5c).

We noted some differences between the two cell lines. The effects of either knockdown were notably stronger and more diverse in H4 cells, illustrated by the lower *R^2^* value produced when correlating DEGs by log_2_FC (Supplementary Fig. 5c). Furthermore, whilst no significant changes in expression of other NSL or Male-Specific Lethal (MSL) complex genes were observed in SH-SY5Ys, upregulation of three genes, namely *HCFC1, WDR5* and *MSL1*, were recorded following *KANSL1* knockdown in H4s, pointing towards the compensatory increase in expression of complex machinery (Supplementary Fig. 5a). These diverging patterns in expression appeared to mirror the distinct differential gene expression patterns observed in glial over neuronal cell types in human brain (Fig. 3).

Next, we characterised the functions of the *KANSL1* DEGs by examining the enriched GO pathways in both cell lines. This analysis identified 365 and 1482 GO terms in SH-SY5Ys and H4s respectively (Fig. 4b, Supplementary Fig. 6). Upregulated DEGs in SH-SY5Ys were enriched for terms such as regulation of gene expression (GO:0010468, FDR range = 3.54 x 10^−13^ – 1.76 x 10^−7^), whilst upregulated DEGs in H4s were enriched for positive regulation of transcription by RNA polymerase II (GO:0045944, FDR range = 3.64 x 10^-23^ – 1.78 x 10^-2^), indicating a compensatory increase in production of transcriptional machinery in response to NSL complex perturbation. Interestingly, amongst the terms enriched within downregulated H4 DEGs were terms relating to mitochondrial function (mitochondrial matrix, GO:0005759, FDR range = 3.39 x 10^-8^ - 2.38 x 10^-2^), autophagy (autophagy, GO: 0006914, FDR range = 4.06 x 10^-6^ – 7.50 x 10^-3^) and lysosomal function (lysosome localisation, GO: 0032418, FDR range = 6.02 x 10^-4^ – 2.43 x 10^-2^), all of which have been implicated in PD pathogenesis (Supplementary Fig. 6)[9]. Given this finding and the fact that prominent lysosomal genes linked to PD risk, including *GBA1* and *CTSB*, were found to be dysregulated following NSL complex perturbation, we examined the enrichment of a broader list of genes linked to the lysosomal system within the two DEG lists (Supplementary Fig. 7a, Supplementary Fig. 8)[46]. Published by Brozzi et al., 2013, this list was generated by collecting and annotating lysosomal gene lists from proteomic studies, databases, reviews and systems biology approaches. We found this list was significantly enriched within downregulated *KANSL1* DEGs in both SH-SY5Ys (FDR = 5.90 x 10^-7^) and H4s (FDR =1.28 x 10^-2^) (Supplementary Fig. 8). Taken together, these results suggest that the NSL complex plays a substantial role in the regulation of lysosomal function.

### *KANSL1* knockdown dysregulates Parkinson’s disease-associated genes and pathways

To investigate the relevance of the genes dysregulated by *KANSL1* in PD, we assessed the enrichment of genes causally linked to Mendelian and sporadic forms of the disease[3,44,45]. In SH-SY5YS, we found that 23 PD-associated genes (nine upregulated, 14 downregulated) were differentially expressed following *KANSL1* knockdown (Fig. 4c, Supplementary Fig. 7a). Using GSEA, we revealed a significant enrichment of genes relating to sporadic PD in these DEGs (FDR = 1.9 x 10^−2^) and a nominally significant enrichment among downregulated DEGs in particular (FDR = 7.91 x 10^-2^)(Fig. 4c, Supplementary Fig. 7a). In H4s, 62 PD-associated genes were dysregulated (23 downregulated, 39 upregulated), although this did not result in a significant enrichment of either gene list.

Next, we assessed the association of PD-relevant pathways within cell lines[47]. In SH-SY5Ys, we detected the enrichment of just one such pathway, namely the Lysosome pathway (KEGG:04142, FDR = 3.59 x 10^−2^), in *KANSL1* DEGs. This remained the sole pathway to be enriched when examining downregulated DEGs (FDR = 1.78 x 10^−2^) separately to upregulated, indicating that these genes in particular are driving this enrichment. This pathway was also found to be enriched within H4 DEGs (FDR = 1.48 x 10^-4^), along with Neutrophil degranulation (FDR = 1.95 x 10^-2^). Together these findings, suggest a role for KANSL1 in lysosome and immune function, both processes that were highlighted in our analyses of post-mortem human brain tissue.

### Direct targets of *KANSL1* regulation are enriched for lysosomal pathways

We noted that *KANSL1* knockdown could impact on gene expression both directly, through its activity within the NSL complex, and indirectly, through secondary effects on cell function. Given that we are most interested in the former, we leveraged publicly available KANSL1 ChIP-sequencing data conducted in HeLa cells[16]. This study detected 3637 genes bound by KANSL1 at transcription start sites across three replicate experiments. Upon filtering for genes differentially expressed following *KANSL1* KD in either cell line, we identified 298 common hits in SH-SY5Ys and 1619 common hits in H4s, both of which denoted a significant enrichment (*p-*value range = 2.2 x 10^-16^ – 1.44 x 10^-10^), with 163 genes reaching significance in all three experiments (Supplementary Fig. 9). Given the pro-transcriptional nature of the NSL complex, we predicted that genes which were downregulated were more likely to be directly regulated by the NSL complex. In SH-SY5Ys, these 163 common hits were enriched for several terms relating to lysosomal function, such as lysosomal organisation (GO:0007040, FDR = 7.62 x 10^-5^), as well as autophagy (GO:0006914, FDR = 7.82 x 10^-3^) (Fig. 4d)[55]. Similar terms were also found to be enriched in the 1004 common hits in H4s, including lysosomal membrane (GO:0005765, FDR range = 7.61 x 10^-3^ – 2.74 x 10^-2^) and autophagy (GO:0006914, FDR range = 7.18 x 10^-3^ – 1.50 x 10^-2^).

Furthermore, we examined the relevance of these common target genes in PD. Within this SH-SY5Y list, we found three relating to sporadic forms of PD, namely *PARP9* (encoding the inflammatory mediator, Poly(ADP-ribose) Polymerase 9), *USP8* (encoding the endosomal trafficking protein, ubiquitin specific protease 8) and *GBF1* (encoding Golgi Brefeldin A Resistant Guanine Nucleotide Exchange Factor 1, which has roles in vesicular trafficking and viral replication), and one Mendelian gene, namely *GBA1*[56–58]. The H4 list contained five different PD-associated genes, namely *NUCKS1, PIGL, RNF141, AREL1* and *SYBU*. Furthermore, upon examining the enrichment of terms implicated in PD through common genetic variation, Lysosome was again the sole pathway found to be enriched within the list of 163 common target genes in SH-SY5Ys (KEGG:04142, FDR = 1.97 x 10^−4^)[47]. Taken together, these findings provide direct experimental support for NSL complex activity at genetic loci associated with PD risk and point towards autophagic and lysosomal mechanisms being particularly important regulatory targets in neuronal and glial cells.

### *KANSL1* contributes to differential gene expression driven by 17q21.31 haplotype variation across major cell types

Finally, we sought to evaluate the contribution of *KANSL1* and the NSL complex to the differential gene expression observed in brain between H2 allele carriers and H1 homozygotes. Given that a significant difference in *KANSL1* expression was detected between haplotype groups most strongly at the level of individual cell types, we focused on the snRNA-sequencing differential gene expression analysis results. As previously mentioned, we predicted that genes downregulated following *KANSL1* knockdown were most likely to represent direct NSL complex targets. Furthermore, we would expect this gene set to have lower expression in H1 homozygotes as compared to H2 allele carriers, assuming that *KANSL1* was indeed a key regulator of their expression (Fig. 5a). In fact, we found that 60.3% and 50.8% of DEGs in SH-SY5Ys and H4s respectively had this pattern of expression in human IPL (Supplementary Fig. 10a,b). Similarly, in ACG these overlaps were 36.8% and 56.6% respectively (Supplementary Fig. 10c,d). Moreover, some of these common DEGs had additional evidence of KANSL1 binding (Fig. 5b, Supplementary Fig. 11a). The dysregulation of *PURG (*which encodes a DNA binding protein) and *WDR48 (*which encodes a deubiquitinase regulator) was observed in H4s as well as IPL astrocytes, whilst *SENP2, USP30* and *GNS* were dysregulated in SH-SY5Ys as well as IPL excitatory neurons[59,60]. *SENP2* encodes an enzyme involved in SUMOylation and mitochondrial modulation, *USP30* encodes a deubiquitinase enzyme which negatively regulates mitophagy, and *GNS* encodes the lysosomal enzyme glucosamine (N-Acetyl)-6- Sulfatase, the deficiency of which leads to a lysosomal storage disorder[61–64]. We further investigated the change in expression of these three genes following *KANSL1* knockdown in iNeurons, which are a more representative human neuronal model, and found that they were indeed downregulated (Supplementary Fig. 12).

**Fig. 5.**
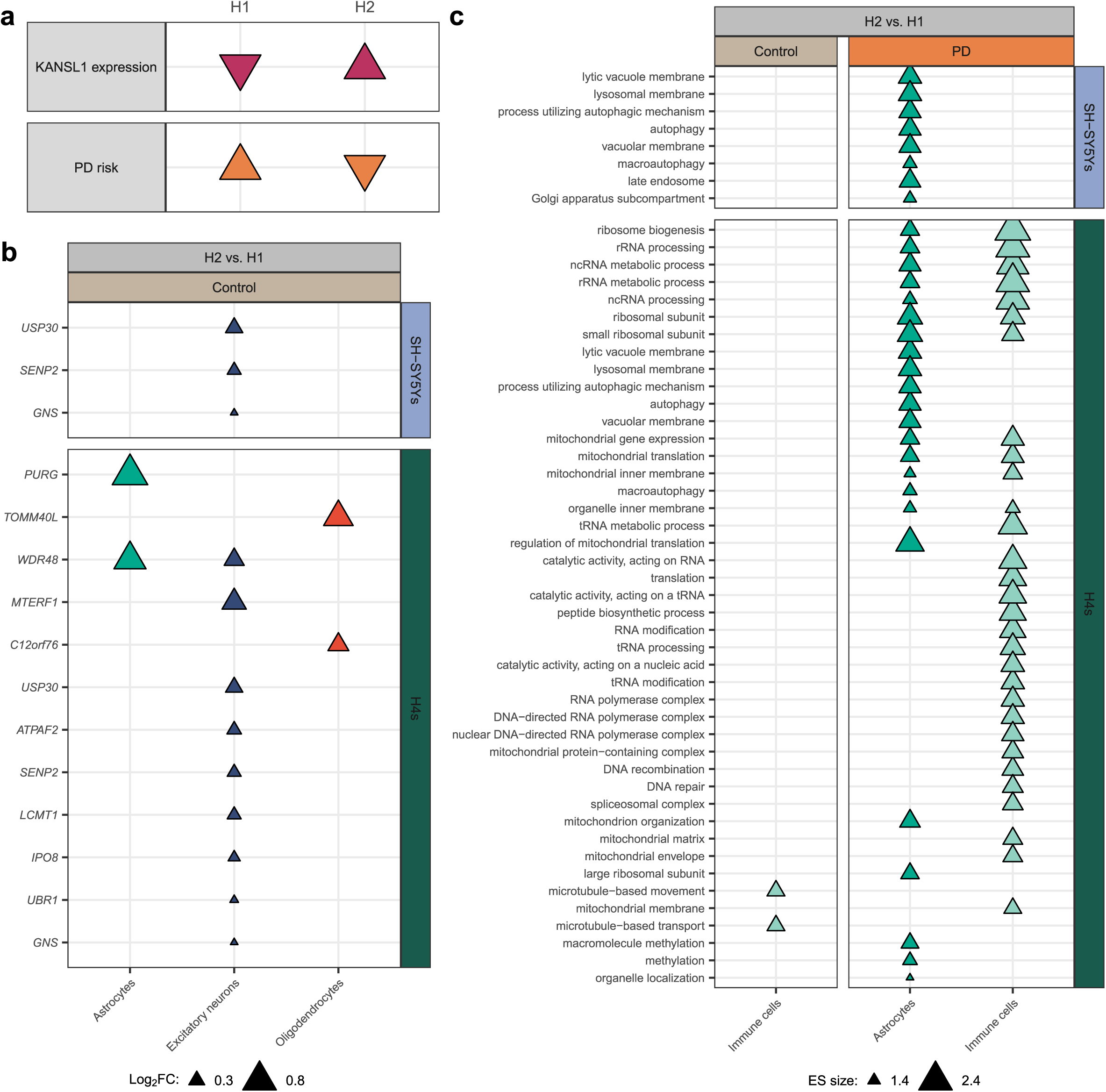
Genes and pathways dysregulated by both 17q21.31 haplotype status in IPL and by *KANSL1* knockdown in cell lines are relevant to PD. A) The 17q21.31 H1 haplotype has been linked to an increase in PD risk and decreased expression of *KANSL1*. B) Common genes that were significantly upregulated in H2 carriers over H1 homozygotes in brain, significantly downregulated following *KANSL1* knockdown in either SH-SY5Ys or H4s, and directly bound by KANSL1 in HeLa cells included genes linked to lysosomal and mitochondrial functions[16]. The log_2_FC displayed is the value of the H2 vs. H1 brain result. C) Common pathways that were enriched among upregulated gene expression in H2 carriers over H1 homozygotes in brain (as determined by ranked GSEA) and among genes both downregulated following *KANSL1* knockdown and directly bound by KANSL1 in HeLa cells included lysosomal and autophagic pathways[16]. Enrichment score (ES) denotes the strength of the enrichment as well as the direction within the rank.

Lastly, we examined the common pathways that were changing in these datasets. Of the GO terms significantly associated with changing gene expression across all cell types in H2 allele carriers compared to H1 homozygotes in IPL, we found 206 terms in common with SH-SY5Y *KANSL1* DEGs and 885 in common with H4 *KANSL1* DEGs. More specifically, when filtering for terms enriched within the two lists of common target genes, we identified eight and 42 terms of interest within SH-SY5Y and H4 gene lists respectively (Fig. 5c). Many of these common terms were also found in ACG (Supplementary Fig. 11b). These terms were associated with dysregulated gene expression in glial cell types and included autophagy (GO:0006914, FDR range = 2.34 x 10^-5^ – 2.77 x 10^-2^) and lysosomal membrane (GO:0005765, FDR = 1.63 x 10^-4^). Taken together, these results provide evidence for the notion that haplotype variation at the 17q21.31 locus results in variation in *KANSL1* expression, which in turn gives rise to small but robust changes in the expression of a range of genes causally implicated in PD, with effects on autophagy and lysosomal function.

## Discussion

To date, there has been no systematic examination of the effect of 17q21.31 haplotype variation on global patterns of gene expression in human brain. To address this knowledge gap, we characterised the effects of the 17q21.31 locus on gene expression in post-mortem human brain, both in bulk tissue and within individual cell types, and specifically assessed the contribution of *KANSL1* using orthogonal, newly generated transcriptomic data from human cell lines. We found that haplotype variation at this locus is associated with differential expression of a range of PD-relevant molecular processes. These include autophagy, lysosomal and mitochondrial pathways, operating in both neuronal and glial cell types, with a net impact on immune responses. We found evidence to suggest that the regulatory effect of the 17q21.31 locus is mediated at least in part through the gene regulatory activity of KANSL1 and the NSL complex of which it is part.

Focusing on our analyses of post-mortem human brain tissue, as we expected, we found strong evidence to support the impact of haplotype variation on the expression of a range of genes within the 17q21.31 locus itself. In fact, of the genes within the locus, we identified 13 that were differentially expressed either using bulk-level or snRNA-sequencing analyses, and results were highly consistent between analyses of control and PD samples. In fact, many of these genes have previously been identified through cis-eQTL analyses using both bulk and, more recently, snRNA-sequencing data derived from post-mortem human brain tissue[12,13]. Importantly, we found that multiple genes previously posited to be drivers of PD risk at the locus, including *LRRC37A/2* and *KANSL1*, were among those differentially expressed[10,18]. However, we noted the absence of data supporting *MAPT,* for which the evidence of differential expression by haplotype has been inconsistent in brain and blood[10,11,65,66]. Interestingly, we found that *KANSL1* was up upregulated across most cell types in individuals carrying the H2 allele, whether in control individuals or in the context of PD. Thus, these analyses suggest that *KANSL1* is one of a number of genes at the 17q21.31 locus which could drive PD risk through changes in RNA expression.

Perhaps more surprisingly, beyond the locus itself, both bulk-level and single nuclear transcriptomic data highlighted a number of molecular processes and biological pathways known to be of key importance in PD. Focusing on the former, we identified significant differences in the expression of genes associated with autophagy, lysosomal and mitochondrial function when stratifying by 17q21.31 haplotype. Critically, we found that the expression of these pathways was higher in control and PD-affected individuals carrying the H2 allele. Since this allele is known to be protective in PD, these findings are highly consistent with existing data suggesting that reduced function of these core cellular processes can drive PD risk in both familial and complex forms of disease[9]. Furthermore, we found that genes and pathways relating to the immune system, including both innate and adaptive responses, were elevated in individuals carrying the H2 allele. This was apparent in control and PD-affected individuals at the bulk-level, with a more complex picture in single cells, where we found that this appeared to be driven by effects on glial cell types, particularly astrocytes. Since autophagy, lysosomal and mitochondrial function have all been implicated in modulation of the immune system, including in the context of PD, it seems likely that the impact of haplotype variation at the 17q21.31 locus on immune function is a system effect arising from relatively small changes in these core cellular processes, primarily in glia[67,68].

While the findings in human brain tissue are of considerable interest, alone they do not address the question of whether changes in *KANSL1* expression could explain the observed differences in cellular processes and immune function. This was therefore the main driver for the generation of orthogonal cellular data. By knocking down *KANSL1* in neuronal and glial cell lines, we modelled the impact of the PD risk-associated H1 haplotype in two relevant cellular contexts. Given that *KANSL1* encodes a component of the chromatin-modifying NSL complex, we expected a knockdown to generate widespread differences in gene expression that would be highly correlated with expression signatures generated through perturbing other key components of the complex. Indeed this was the case, with *KANSL1* knockdowns in both cell lines resulting in high numbers of DEGs that significantly correlated with gene expression patterns detected following equivalent *KAT8* knockdowns. Interestingly, *KANSL1* knockdown altered the expression of a significant number of genes causally-associated with PD through genome-wide association studies[2,45]. Whilst we recognise that there is uncertainty over the precise risk-associated genes, several genes found here to be differentially regulated by *KANSL1* are among those with the strongest evidence of a disease association, including *GBA1* and *GCH1*[69–71]. Similarly, *PINK1* dysregulation was detected in both cell types following *KANSL1* knockdown, validating a previous finding and connecting *KANSL1*-dependent regulatory processes to familial forms of PD[18]. However, most importantly, the analyses conducted in both cell types highlighted the role of KANSL1 and the NSL complex in regulating autophagy, lysosomal and mitochondrial processes, not only in neurons but also in glia, and with clear implications for PD risk. While these analyses do not rule out contributions from other genes within the 17q21.31 locus in modifying pathway activity in human brain, these results are highly suggestive, given the known role of KANSL1 in gene expression regulation through the NSL complex as well as supporting evidence provided by public ChIP-sequencing data.

Nonetheless, we recognise that further analyses are required to fully understand the role of *KANSL1* and the 17q21.31 locus in PD. We note that all the brain samples used in this analysis were derived from post-mortem human tissue and, in the case of PD-affected individuals, focused on the late stages of disease. Given that across the whole lifetime of an individual, compensatory processes may be triggered to mask the potential effects of altering gene expression levels within the locus, it would be beneficial to study the impact of genetic variation therein across multiple disease stages and time points, together with data from other human-derived tissues. Furthermore, while cell lines such as SH-SY5Ys and H4s allow for rapid genetic manipulations in disease-relevant backgrounds, these models may not fully recapitulate the processes of disease-affected tissues, and may be particularly limited when attempting to study the role of the immune system. Finally, and perhaps most importantly, this work raises the need for generating additional regulatory data at a cell type-specific level, including further information on chromatin structure and organisation in relation to KANSL1-dependent NSL complex activity.

Despite these limitations, this study significantly advances our understanding of the 17q21.21 locus and specifically the role of *KANSL1* in PD. We demonstrated that the H1 PD risk haplotype dysregulates the autophagy-lysosome system, potentially leading to immunological changes primarily conferred by glial cell types. Thus, we highlight links between NSL complex activity and PD aetiology that may represent therapeutically viable targets.

## Data availability

Bulk and snRNA-sequencing data derived from human brain can be accessed via the CRN cloud here: <u>10.5281/zenodo.14373344</u> and <u>10.5281/zenodo.14373048</u>. Code used to analyse these two datasets will be made available upon publication here: https://github.com/amyrosehicks/RNAseq_17q21_brain.git and https://github.com/amyrosehicks/snRNAseq_17q21_brain.git. Bulk RNA-sequencing data from SH-SY5Y and H4 cell samples can be accessed from Gene Expression Omnibus using this accession code: GSE283747. Code used to analyse this data will be made available upon publication here: https://github.com/amyrosehicks/RNAseq_K1K8kd.git.

## Acknowledgements

A.H. was supported through the award of an Eisai-Leonard Wolfson Doctoral Training programme in Neurodegeneration. This research was funded in part by Aligning Science Across Parkinson’s (grant number ASAP 000478) through the Michael J. Fox Foundation for Parkinson’s Research (MJFF) as well as MJFF grant number 023434. For the purpose of open access, the author has applied a CC BY public copyright license to all Author Accepted Manuscripts arising from this submission. RHR is currently employed by CoSyne Therapeutics (Lead Computational Biologist), all work performed for this publication was performed in her own time, and not as part of her duties as an employee.

## Supplementary figure legends

**Supplementary** Fig. 1 **Enrichment of GO terms across changing gene expression at bulk tissue level in three clinical groups, as determined by ranked GSEA.** Child terms enriched across gene expression in both directions were excluded. The terms displayed are parent terms, with the number of reduced child terms displayed in brackets (term reduction utilised a hierarchical tree cut off threshold of 0.9). Terms are ranked by their prevalence across clinical groups and by the strength of the enrichment score (ES).

**Supplementary** Fig. 2 **The 17q21.31 locus drives cell type-specific patterns of differential gene expression in ACG.** A) DEGs were detected across nearly all cell types in both control and PD samples when comparing H2 allele carriers to H1 homozygotes. B) The majority of differential gene expression detected in this analysis were cell type-specific, in both upregulated and C) downregulated directions. D) We detected differential expression of 10 genes within the 17q21.31 locus, three of which were specific to one cell type in one clinical group, with the remaining seven dysregulated across multiple cell types. E) Using ranked GSEA, we detected the enrichment of GO pathways across differential gene expression in all major cell types, particularly in astrocytes and immune cells. Enrichment score (ES) denotes the strength of the pathway enrichment as well as the direction within the ranked results. F) Pathways genetically-linked to PD through polygenic risk scoring were also found to be enriched across changing gene expression in five major cell types using ranked GSEA[47].

**Supplementary** Fig. 3 **Enrichment of GO terms across changing gene expression in IPL control samples and PD samples and across major cell types, as determined by ranked GSEA.** The terms denote parent terms, with the number of reduced child terms displayed in brackets. Term reduction utilised a hierarchical tree cut off threshold of 0.9. Child terms enriched across gene expression in both directions in the same cell type are excluded from the plot.

**Supplementary** Fig. 4 **Enrichment of GO terms across changing gene expression in ACG control samples and PD samples and across major cell types, as determined by ranked GSEA.** The terms denote parent terms, with the number of reduced child terms displayed in brackets. Term reduction utilised a hierarchical tree cut off threshold of 0.9. Child terms enriched across gene expression in both directions in the same cell type are excluded from the plot.

**Supplementary** Fig. 5 **NSL complex knockdown in SH-SY5Ys and H4s.** A) NSL and MSL complex genes significantly differentially expressed following either *KANSL1* or *KAT8* knockdown in H4 and SH-SY5Y cells. B) Total number of differentially expressed genes following *KAT8* knockdown detected across both cell lines. C) Correlation in log_2_FC in gene expression following either *KANSL1* or *KAT8* knockdown, across both cell lines. Genes are separated into those differentially expressed in either knockdown (or both) and those not reaching significance. X=Y is displayed on the graph for reference as a dashed line.

**Supplementary** Fig. 6 **Enrichment of GO terms among *KANSL1* DEGs.** A) Enrichment of SH-SY5Y DEGs. B) Enrichment of H4 DEGs (filtered for molecular function and cellular component terms). C) Enrichment of H4 DEGs (filtered for biological process terms). Parent terms are displayed, with the number of reduced child terms displayed in brackets (term reduction utilised a hierarchical tree cut off threshold of 0.7). Triangle size corresponds to the ratio (intersection size/term size) of the child term with the highest significant FDR. Child terms enriched across gene expression in both directions in the same cell type are excluded from the plot.

**Supplementary** Fig. 7 **PD-associated differential gene expression in SH-SY5Ys and H4s following *KANSL1* knockdown.** A) Differential expression of PD-associated genes across both cell lines following *KANSL1* knockdown, with DEGs common to both cell types (displayed in Fig. 4c) excluded. B) Association of PD-related gene lists with changing gene expression following KANSL1 knockdown in both cell lines, as determined by ranked GSEA. Enrichment score (ES) denotes the strength of the association[2,44,45].

Supplementary Fig. 8 Differential expression of genes associated with lysosomal function across SH-SY5Ys and H4s following *KANSL1* knockdown. Genes are ordered by log_2_FC and split into two columns for visualisation purposes[46].

**Supplementary** Fig. 9 **Overlap between our RNA-sequencing results and published ChIP-sequencing results**. Plot displays overlaps between genes identified as differentially expressed following *KANSL1* knockdown in SH-SY5Y cells or H4 cells, and genes found to be directly bound by KANSL1 through ChIP-sequencing of HeLa cells[16].

**Supplementary** Fig. 10 **Overlap in DE results between brain bulk and cell line snRNA-sequencing results.** A) Overlap between DEGs identified in IPL (summed across cell types) and SH-SY5Ys. B) Overlap between DEGs identified in IPL (summed across cell types) and H4s. C) Overlap between DEGs identified in ACG (summed across cell types) and SH-SY5Ys. D) Overlap between DEGs identified in ACG (summed across cell types) and H4s.

**Supplementary Fig. 11 Overlap in genes and pathways dysregulated by haplotype status in ACG and by *KANSL1* knockdown in SH-SY5Y and H4 cells.** A) Several genes were significantly upregulated in H2 carriers over H1 homozygotes in brain, significantly downregulated following *KANSL1* knockdown in either SH-SY5Ys or H4s, and directly bound by KANSL1 in HeLa cells[16]. The log_2_FC displayed is the value of the H2 vs. H1 brain result. B) Common pathways that were enriched among upregulated gene expression in H2 carriers over H1 homozygotes in brain (as determined by ranked GSEA) and among genes both downregulated following *KANSL1* knockdown and directly bound by KANSL1 in HeLa cells included autophagy-related pathways[16]. Enrichment score (ES) denotes the strength of the enrichment as well as the direction within the rank.

**Supplementary** Fig. 12 **RT-qPCR of target genes following lentiviral knockdown of *KANSL1* in iNeurons.** A) *KANSL1*, B) *USP30*, C) *SENP2,* and D) *GNS* expression were measured in iNeurons following *KANSL1* knockdown. These genes were examined specifically due to evidence of them being regulated by 17q21.31 locus specifically through KANSL1 activity (Fig. 5b).

