## Supplementary figures and images for "17q21.31 locus regulates Parkinson’s disease relevant pathways through *KANSL1* activity"

### Supplementary Fig. 1

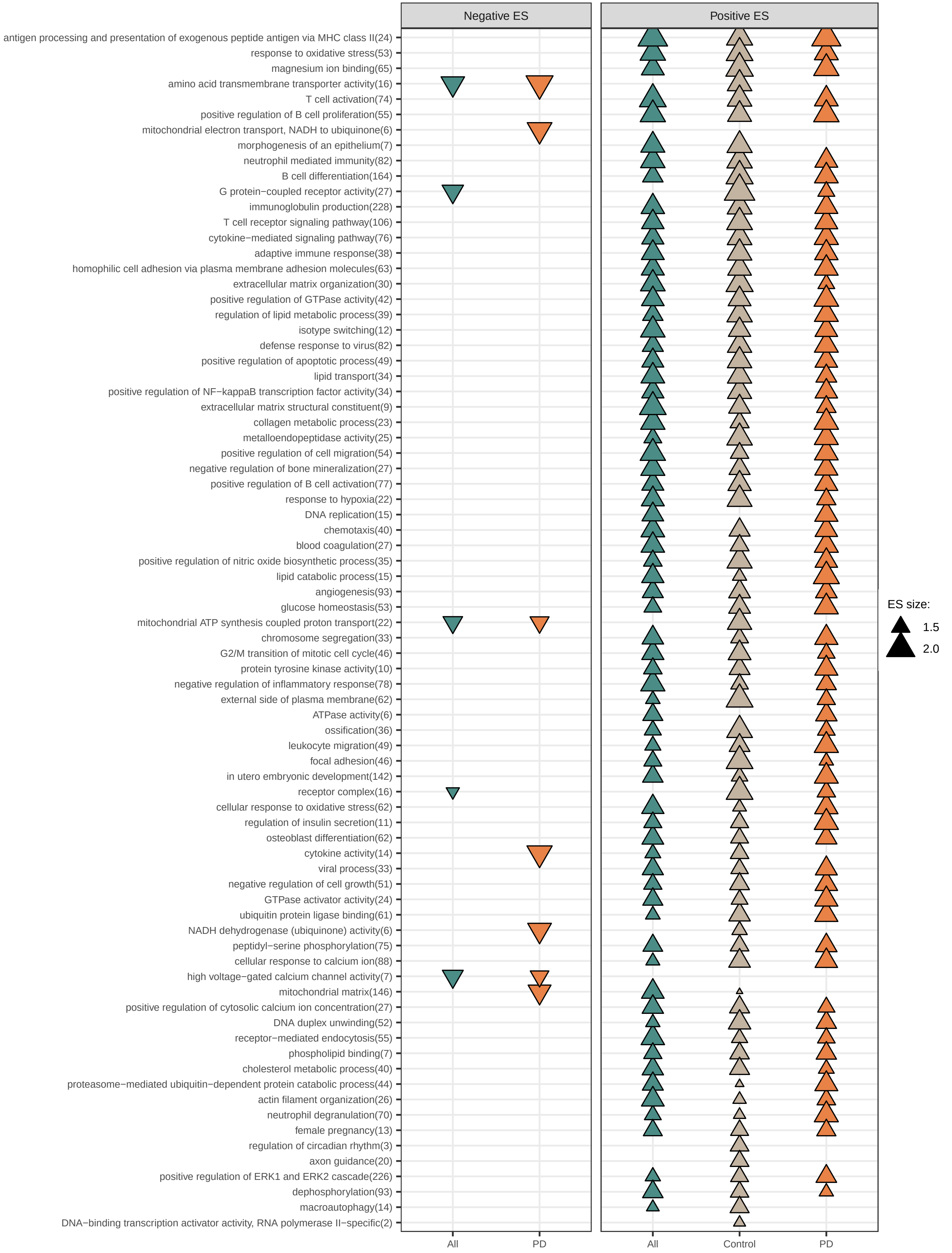

### Supplementary Fig. 2

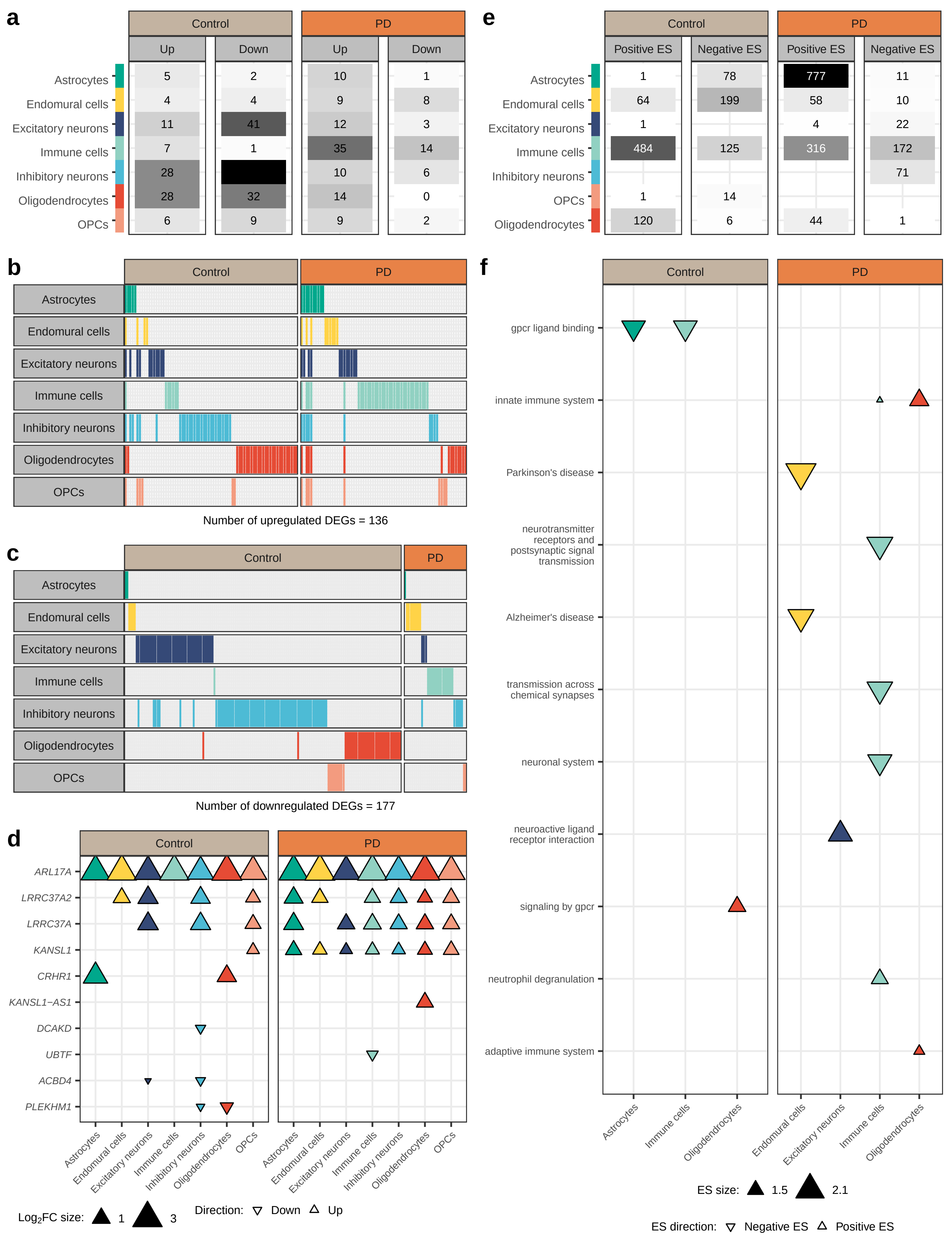

### Supplementary Fig. 3

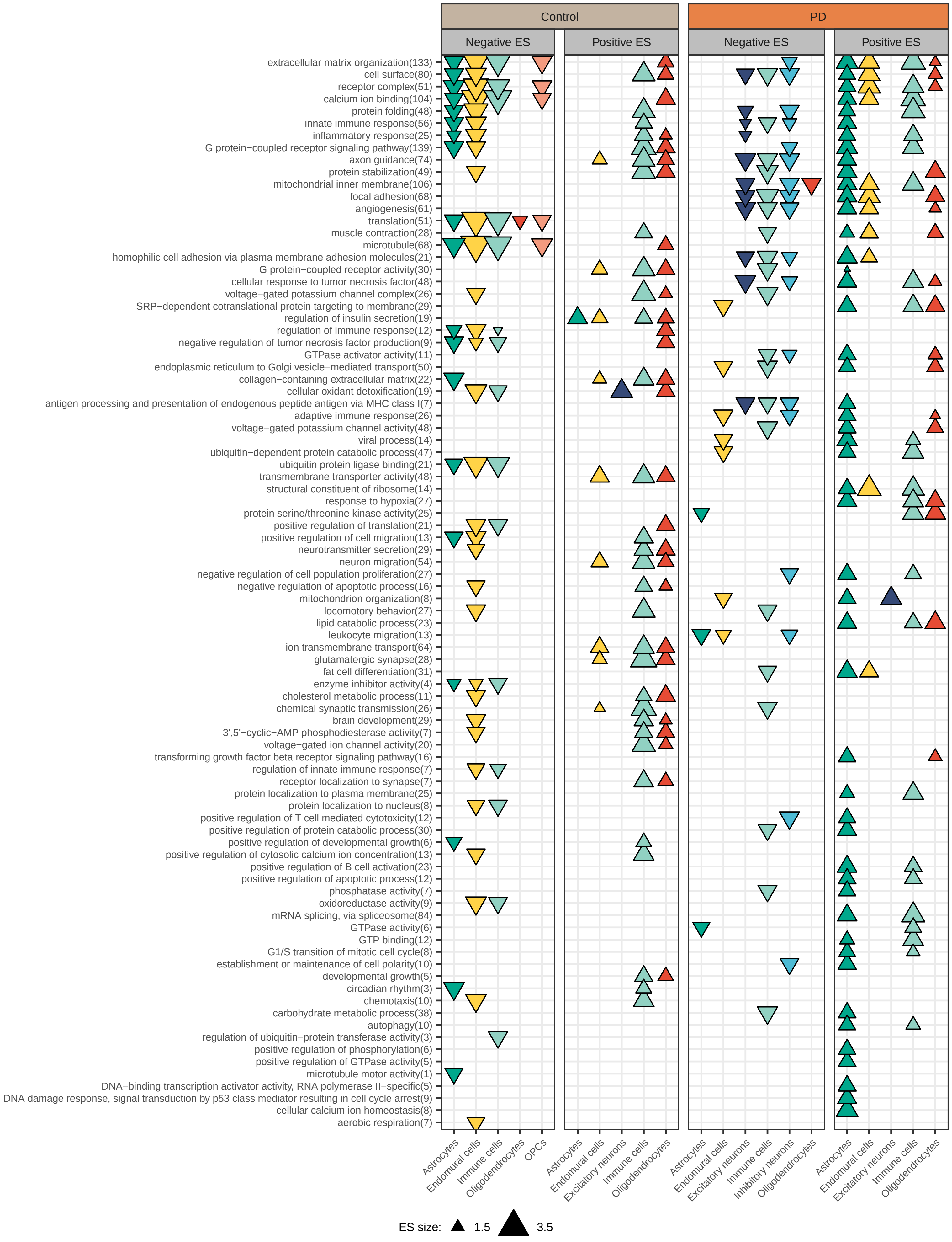

### Supplementary Fig. 4

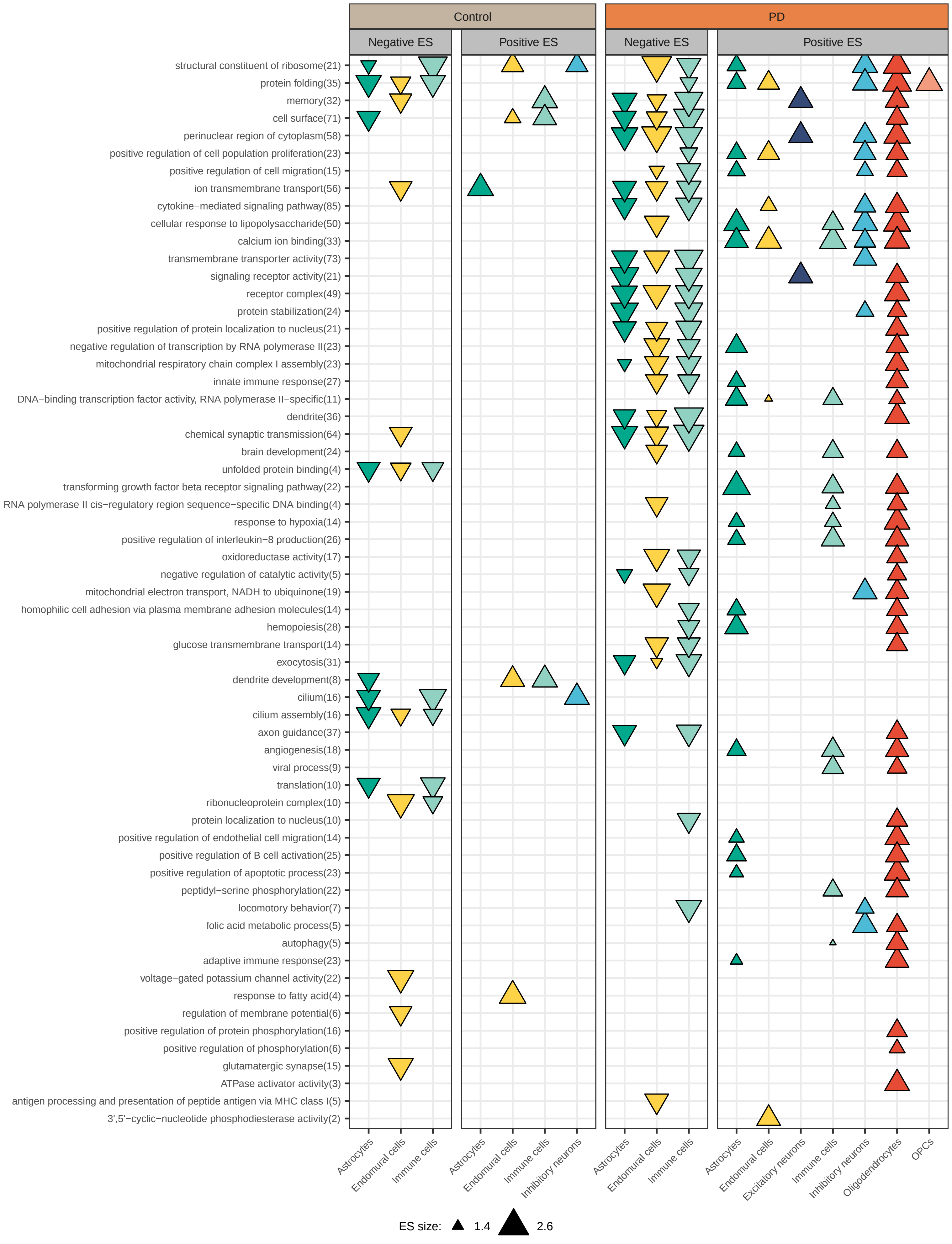

### Supplementary Fig. 5

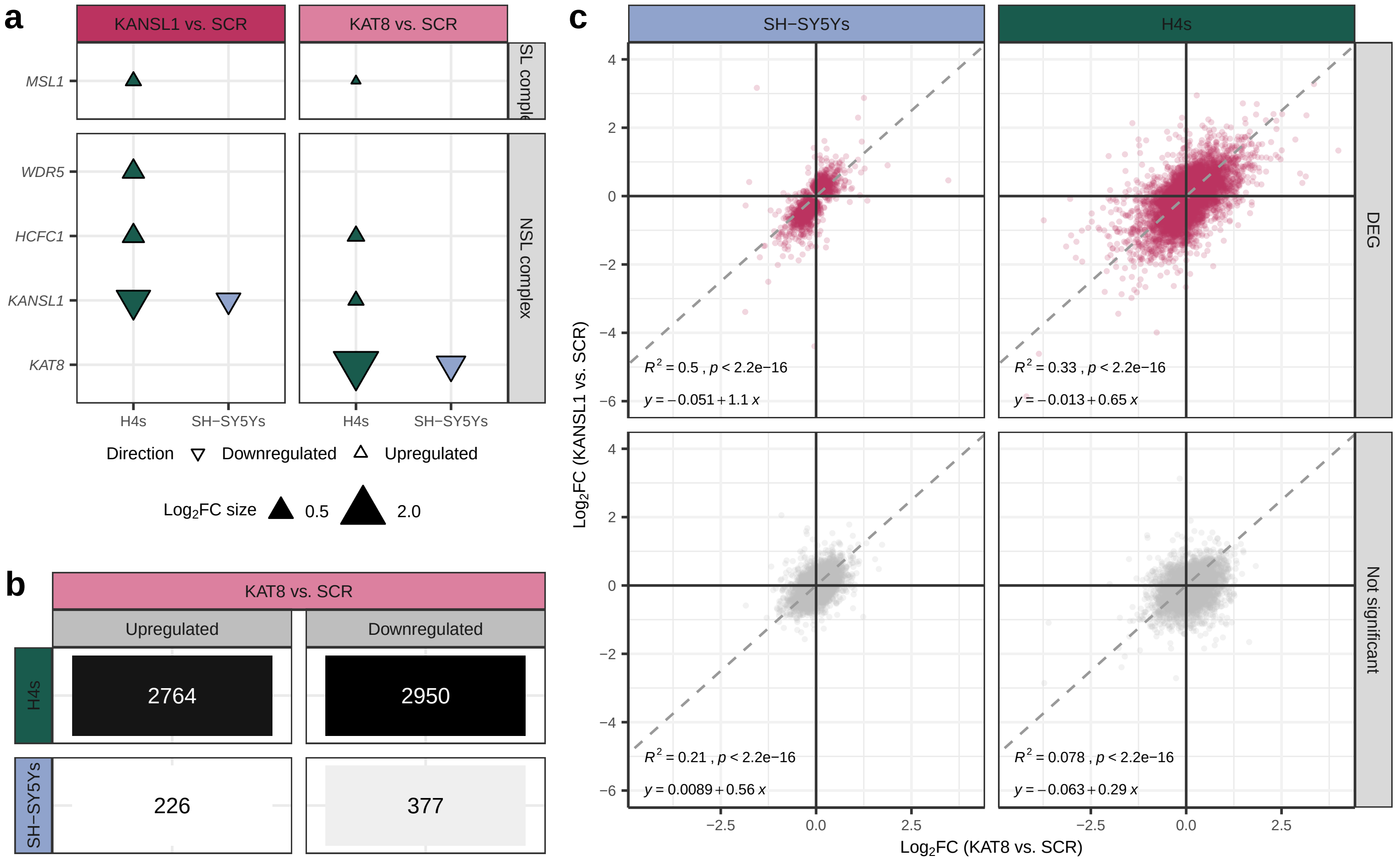

### Supplementary Fig. 6

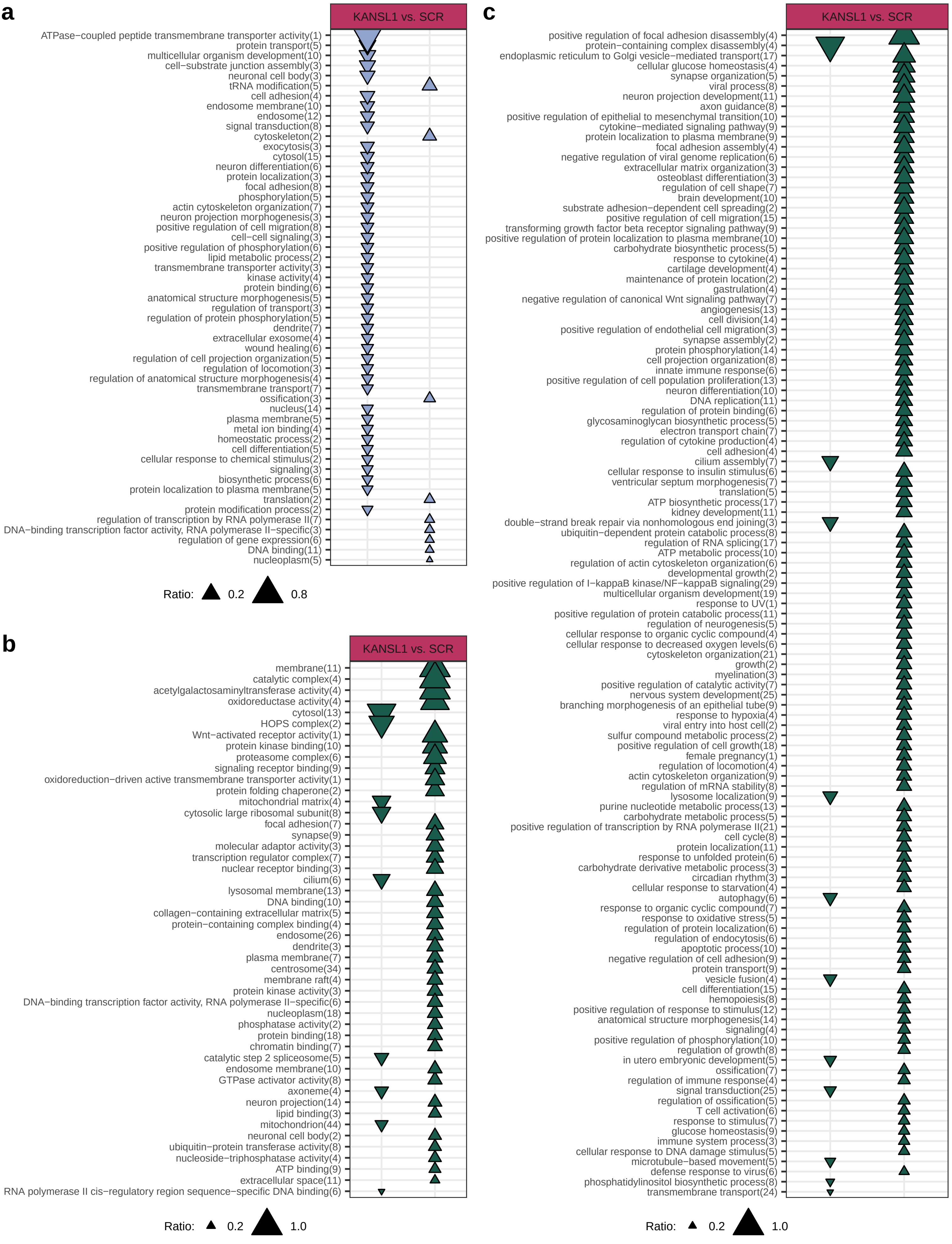

### Supplementary Fig. 7

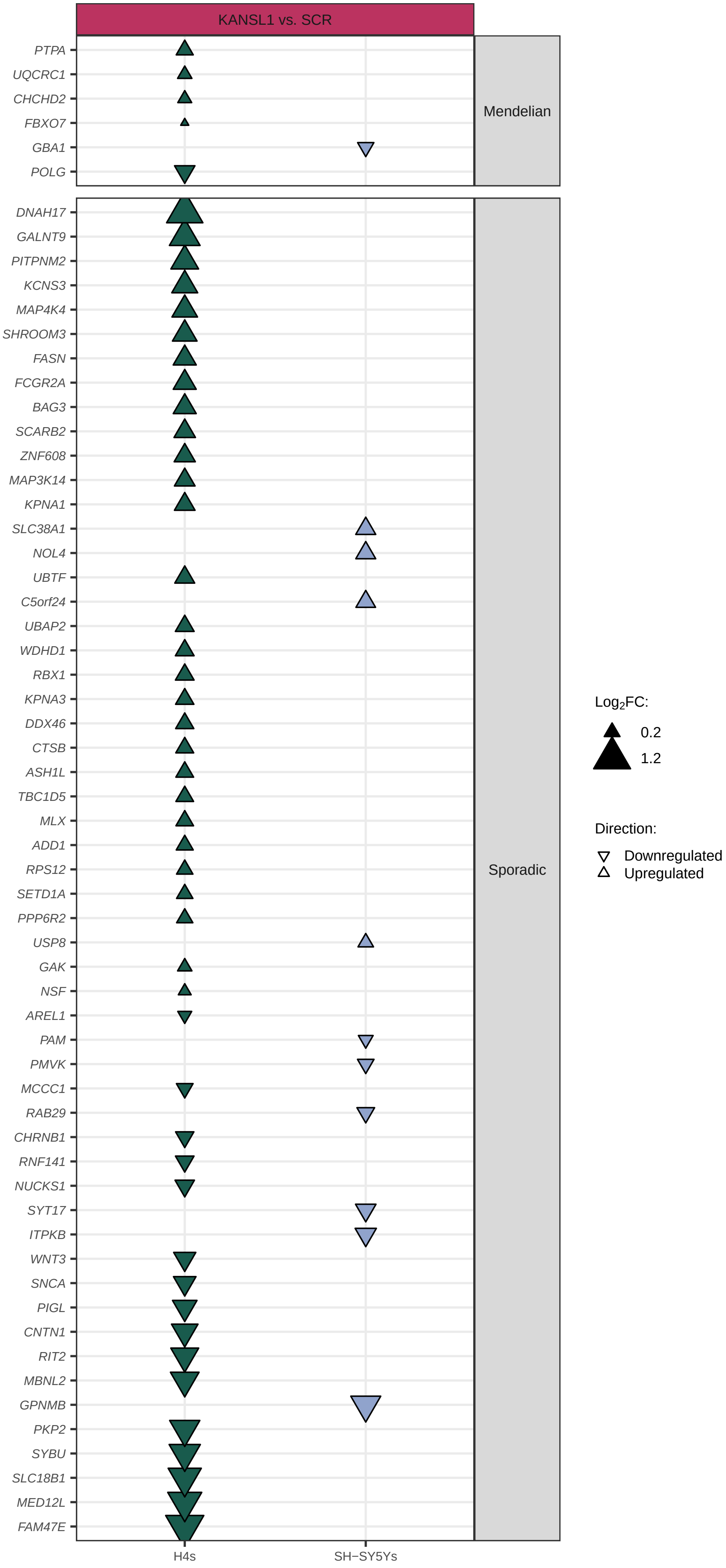

### Supplementary Fig. 8

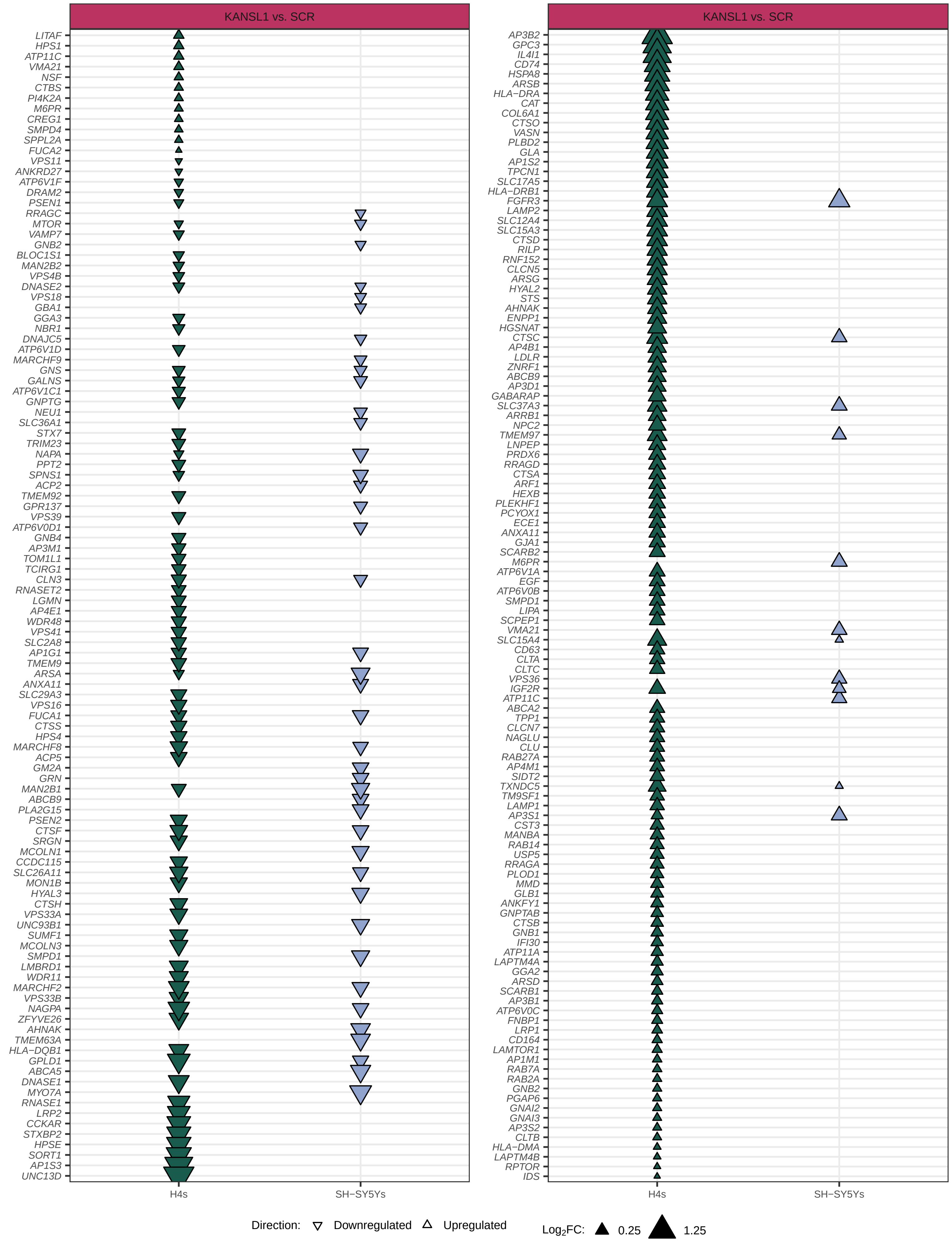

### Supplementary Fig. 9

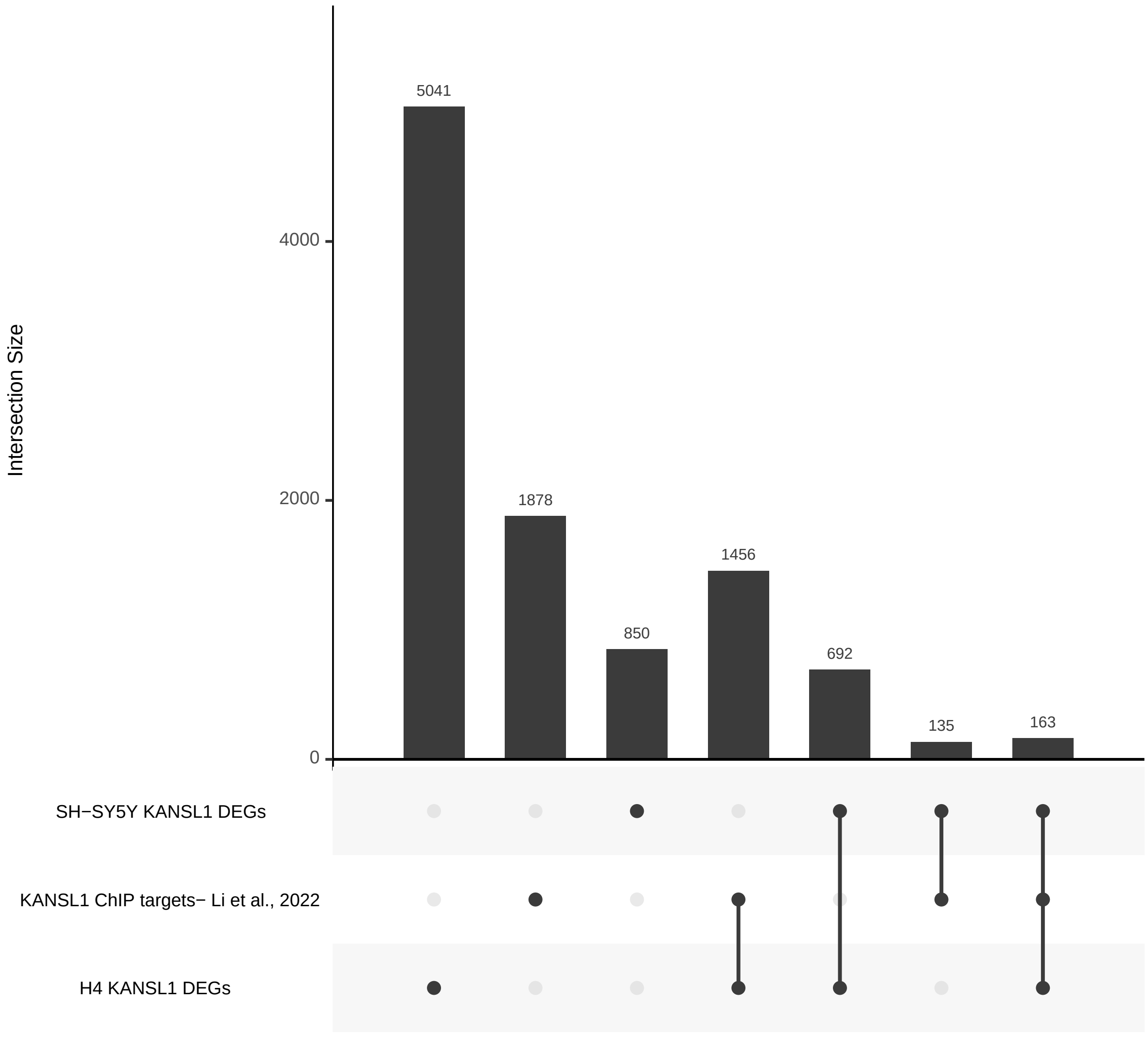

### Supplementary Fig. 10

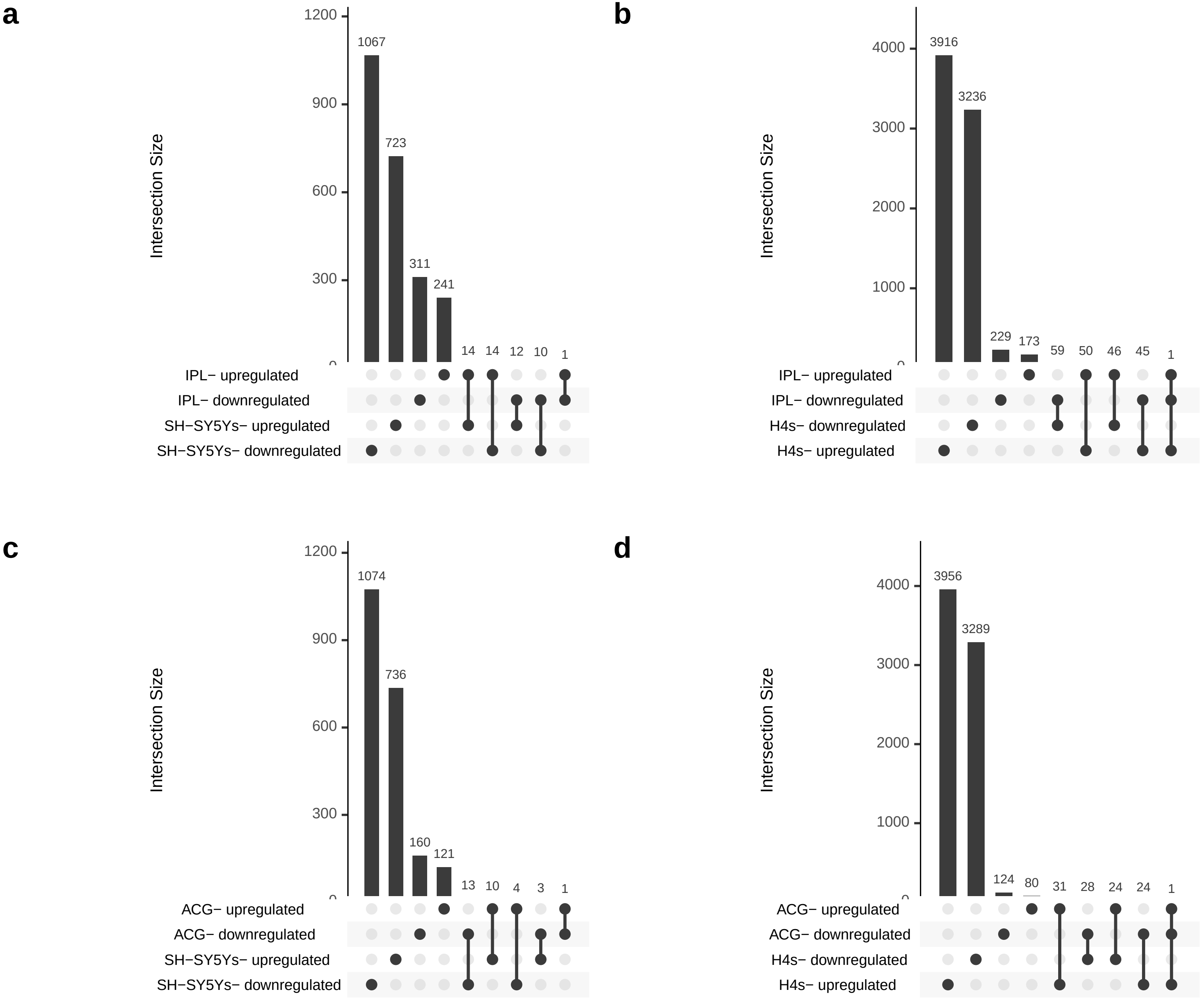

### Supplementary Fig. 11

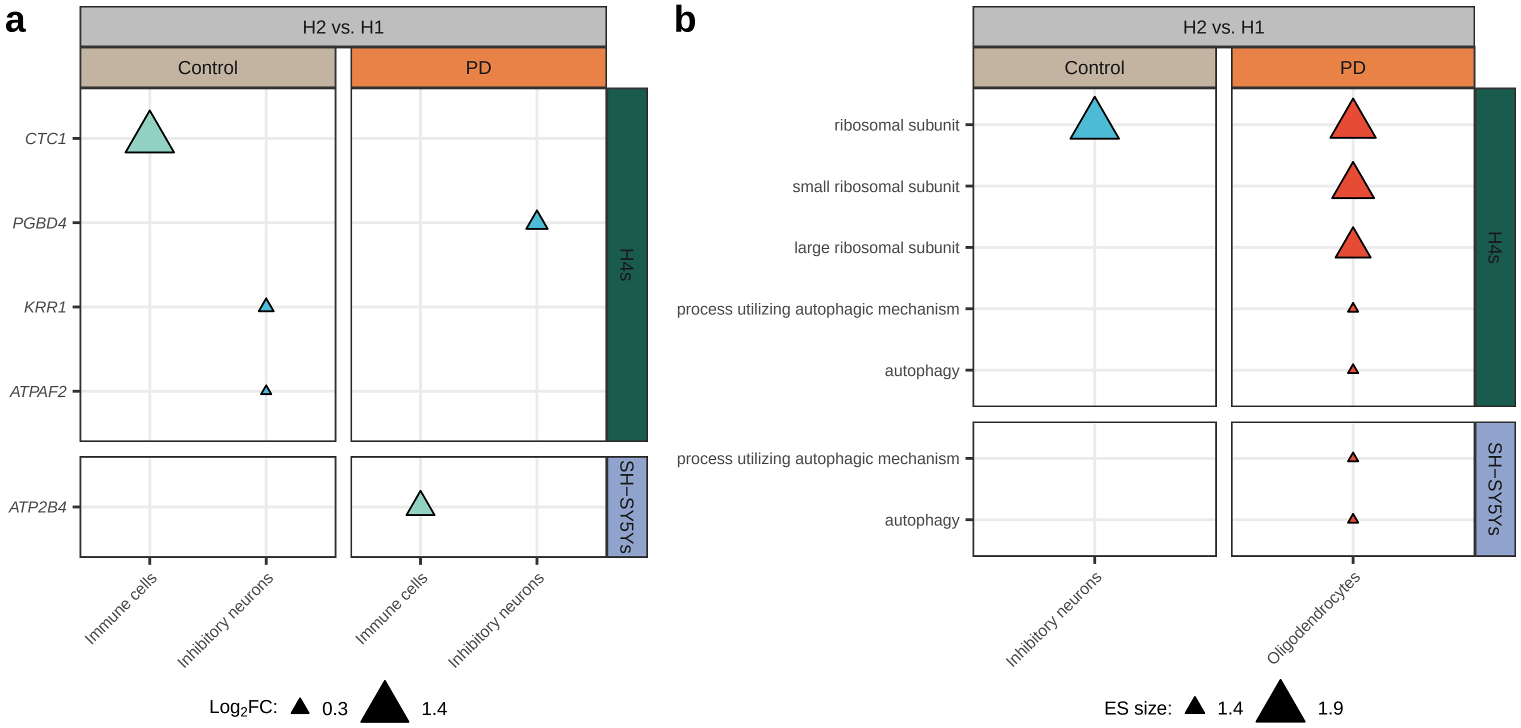

### Supplementary Fig. 12

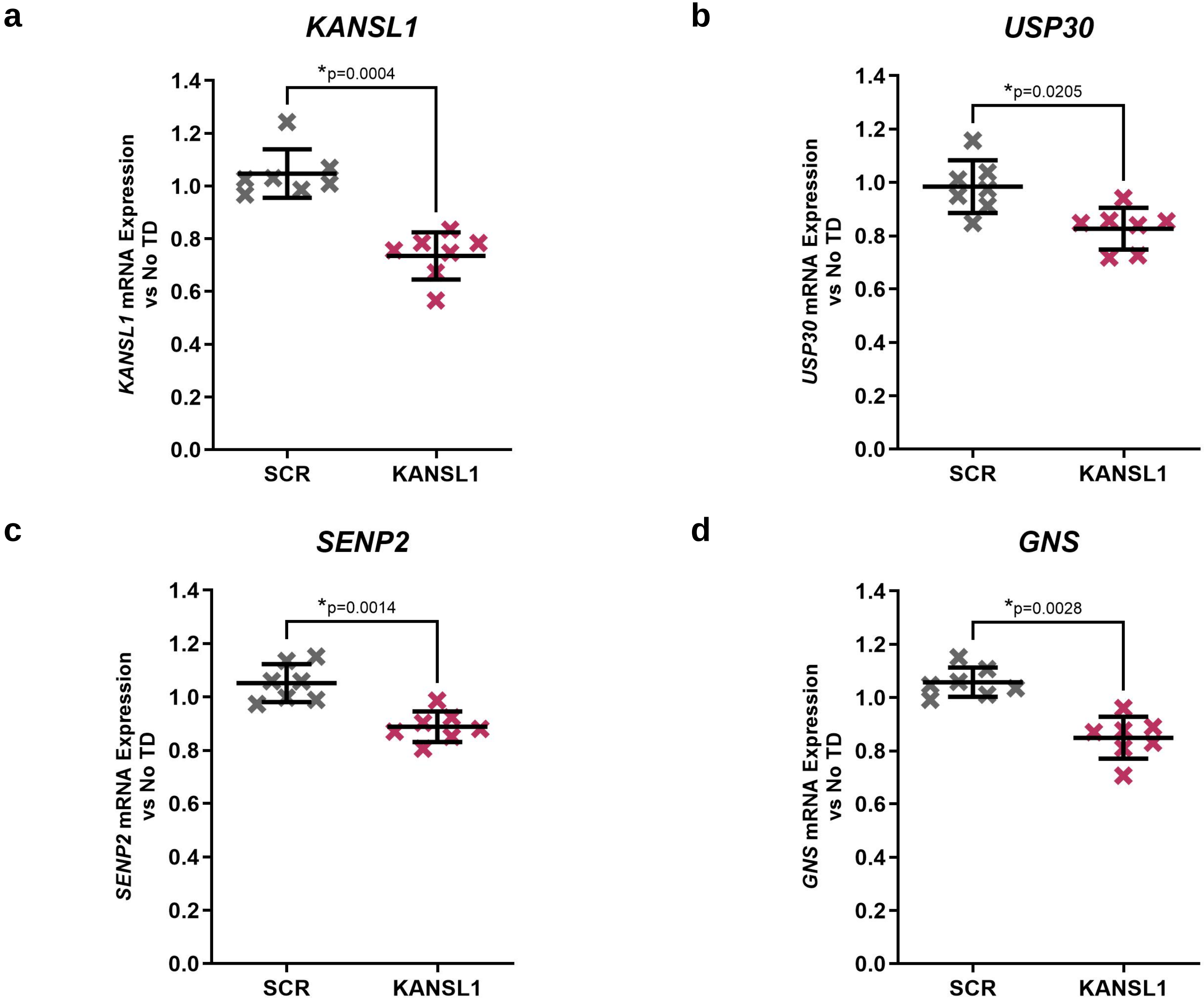
